# Path-dependent recovery of the gut microbiome after antibiotics emerges from coupled ecological and evolutionary dynamics

**DOI:** 10.64898/2026.05.22.727306

**Authors:** Chinmay Lalgudi, Masakazu Kotaka, Eitan Yaffe, Jamie A. Lopez, Feiqiao Brian Yu, Katharine Ng, Justin L. Sonnenburg, Benjamin H. Good, Kerwyn Casey Huang, Handuo Shi

## Abstract

Recovery of the gut microbiome after antibiotic exposure is often incomplete and variable, and the processes underlying this variation remain unclear. We performed longitudinal shotgun metagenomic sequencing of 2876 daily fecal samples from replicated humanized and conventional mouse cohorts exposed to controlled antibiotic perturbations. Metagenomic profiling recapitulated ecological trajectories previously observed by 16S sequencing, while revealing extensive strain-level dynamics, including reproducible sweeps of standing variants and de novo mutations in antibiotic target sites and regulatory loci. We also identified genetic changes whose effects depended on community composition, competitive release, and perturbation history. Cross-housing experiments revealed bidirectional strain transfer, with antibiotic-induced niche clearance enabling replacement of resident strains. In parallel, phage dynamics were heterogeneous and clustered by cage. Together, these findings show that post-antibiotic microbiome recovery is a path-dependent process shaped by selection, transmission, and phage activity, producing divergent outcomes even among closely matched communities exposed to the same perturbations.

## INTRODUCTION

The gut microbiota performs essential functions for host health, including immune modulation, nutrient provision, and pathogen control^1–5^. Antibiotics, while life-saving, can impose strong selective pressures on this ecosystem, disrupting community composition and function and enabling expansion of opportunistic pathogens^5,6^. Recovery following antibiotic perturbation is often slow and incomplete^7,8^. Although microbial communities can rapidly reorganize into alternative stable states after treatment, their return to the pre-perturbation state may take months and remain incomplete^9–11^. Community restoration can be accelerated by the introduction of strains via fecal microbiota transplant^12,13^. The speed and completeness of recovery vary substantially across individuals^10,11^ and, in mice, are shaped by community composition, diet, and access to environmental strain reservoirs^14^. Moreover, even when species recover, the genomes of surviving populations may be altered, harboring genetic changes that alter antibiotic resistance^7,15,16^, metabolic capacity^17^, or interactions with mobile genetic elements^18^. In particular, temperate phages can further reshape post-antibiotic communities through induction-driven bacterial lysis and strain-level turnover even in the absence of species loss^19,20^. These observations underscore the need for high-resolution approaches to capture the eco-evolutionary and mobilome processes that shape post-antibiotic microbiome reassembly. Despite growing recognition of these factors, it remains unclear how context-dependent selection, microbial transmission, and phage dynamics together determine microbiome recovery trajectories.

Longitudinal metagenomics studies in humans and gnotobiotic mouse models have demonstrated that both evolutionary adaptation and ecological strain transmission play central roles in post-antibiotic microbiome reassembly. In a doxycycline-treated human, rapid sweeps of pre-existing variants occurred even in the absence of detectable changes in species-level abundance^15^. During ciprofloxacin treatment in humans, *de novo* mutations in the DNA gyrase subunit A gene (*gyrA*) can sweep within days and persist for weeks after treatment without measurable fitness cost^16^. In mice, singly housed animals exhibit impaired recovery relative to co-housed counterparts^14^ and increased susceptibility to pathogen invasion^21^, highlighting the importance of environmental reservoirs and transmission for recovery. Strain transfer has also been observed among cohabiting adult humans following antibiotic exposure^10^. Beyond selection on bacterial genomes, antibiotics can also induce prophages and mobilize genetic elements^18,20^, potentially amplifying community disruption and accelerating population turnover. Together, these studies establish that antibiotic recovery reflects the interplay of within-host evolution, between-host transmission, and mobile genetic activity. However, in uncontrolled or sparsely sampled systems, it remains difficult to quantify the relative contributions of these processes or to distinguish reproducible responses from stochastic variation in recovery trajectories.

Gnotobiotic mouse models provide a powerful experimental framework for addressing these challenges because they enable controlled perturbations in near-identical cohorts^22–26^. In a previous study, we used this framework with dense longitudinal 16S rRNA profiling to show that post-antibiotic recovery depends on diet and environmental reservoirs^14^. However, because 16S profiling primarily resolves communities at the species level, the underlying strain-level evolutionary, transmission, and phage dynamics remained unclear. Recent longitudinal metagenomic studies in humans have enabled strain-resolved tracking of microbial populations in longitudinal human studies, revealing within-host adaptation, strain replacement, and transmission^10,15,16,27^. Here, we generate and analyze shotgun metagenomic data from 2876 fecal samples collected in Ng et al.^14^ Focusing on humanized gnotobiotic mice, we show that metagenomic profiling recapitulates species-level ecological dynamics inferred from 16S data while enabling strain-resolved analysis of antibiotic selection, strain transmission, and phage activity. We identify widespread genetic sweeps arising from both standing variation and *de novo* mutations, distinguish recovery driven by regrowth of resident strains from replacement by environmentally transmitted strains, and uncover cage-specific prophage dynamics. Together, these results reveal the strain-level and phage-associated processes that govern microbiome recovery following antibiotic perturbation, providing a mechanistic basis for the divergent trajectories observed across similar communities.

## RESULTS

### Shotgun metagenomics of antibiotic-perturbed humanized mouse microbiota recapitulates 16S-derived ecological dynamics

We previously carried out a set of mouse experiments with dense longitudinal stool sampling during and after antibiotic treatment^14^. In these experiments, ex-germ-free mice colonized with human feces (“humanized”) or mice harboring their native, mouse-adapted gut microbiota (“conventional”) underwent one or two courses of ciprofloxacin or streptomycin treatment (**Table S1**). Ciprofloxacin treatment was additionally coupled to dietary perturbations (standard versus MAC-deficient diet; **Table S1**) and to different housing conditions (singly versus co-housed; **Table S1**). Prior 16S rRNA profiling of these experiments showed that the gut microbiota generally recovered following antibiotic exposure, although recovery trajectories were microbiota- and antibiotic-dependent and were modulated by dietary fiber availability and bacterial transmission from the environment^14^. We therefore reasoned that this experimental framework would provide an ideal setting to investigate strain-level responses to antibiotic perturbations that could lead to path-dependent recovery (**Figure 1A**). To do so, we developed a custom low-volume library preparation protocol for metagenomic sequencing (**Methods**)^28^ and sequenced 2876 mouse fecal samples, yielding a median depth of ∼14 million paired-end reads per sample (45 billion paired reads total).

**Figure 1:**
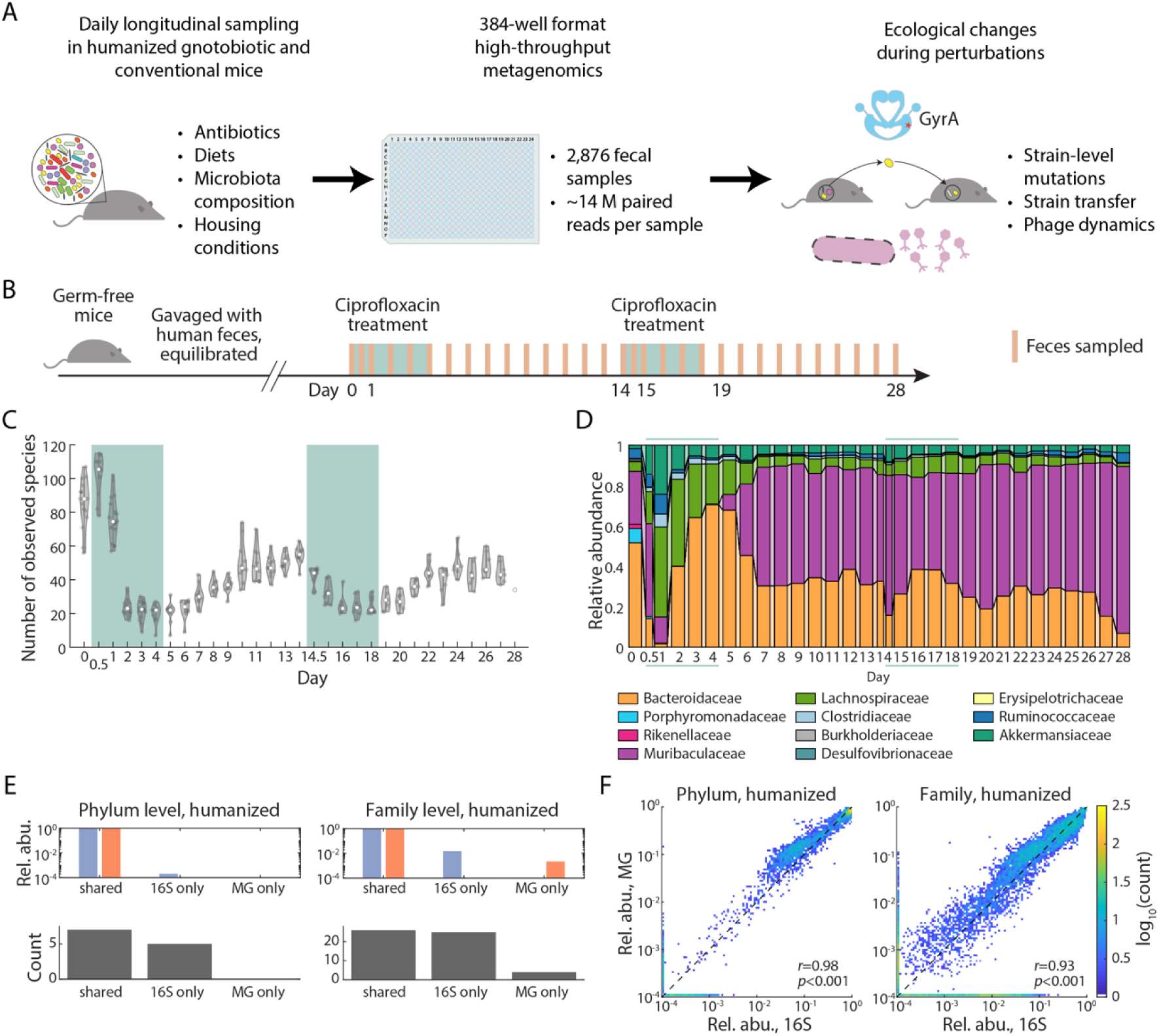
Shotgun metagenomics captures antibiotic-induced ecological restructuring in the gut microbiota of humanized mice. (A) A series of mouse experiments systematically profiling the mouse gut microbiota under perturbations. (B) Experiment schematics for double ciprofloxacin treatment of humanized mice. (C,D) Species-level alpha diversity (richness) (C) and family-level composition (D) changes during the experiment. After the first treatment, the microbiota restructured to a new equilibrium state, and the second treatment had a smaller impact. (E) Phylum-(left) and family-(right) level compositions as detected by 16S and metagenomics in all humanized mice. The two sequencing modalities are consistent at capturing the high-abundance taxa. (F) Heatmaps of abundances at the phylum-(left) and family-(right) levels for all taxa. Metagenomics and 16S are consistent at estimating the relative abundances of all taxa in humanized mice. MG: metagenomics. See also Figure S1.

To determine whether the metagenomics data recapitulated the ecological trajectories previously described by 16S data, we initially analyzed a two-course ciprofloxacin experiment in humanized mice (**Figure 1B**). In this experiment, ten germ-free mice were colonized with feces from a single human donor, allowed to equilibrate for several weeks, treated with ciprofloxacin for five days, allowed to recover for nine days, and then subjected to a second five-day treatment with subsequent recovery (**Methods**). Daily fecal samples were profiled using MIDAS2^29^ with the UHGG database^30^ (**Methods**).

Consistent with prior 16S results^14^, metagenomic profiling revealed a rapid loss of alpha diversity (species richness) during the first ciprofloxacin treatment, followed by partial recovery, with a substantially attenuated effect during the second treatment (**Figure 1C**). Family-level compositional profiles similarly indicated an attenuated response to the second antibiotic exposure, potentially reflecting incomplete recovery prior to re-exposure (**Figure 1D**).

We then extended this comparison across all humanized mouse experiments (87 mice, 1502 fecal samples; **Table S1**) and directly compared metagenomics and 16S sequencing (**Methods**). At the phylum level and family levels, metagenomics captured 7 phyla and 26 families accounting for 99.98% and 98.38%, respectively, of the abundance detected by 16S (**Figure 1E**). The remaining 5 phyla and 25 families were detected by 16S but not by metagenomics. Four additional families were identified by metagenomics but not by 16S; these families accounted for only 0.01%. Thus, the missing taxa from metagenomics account for very little total abundance and are likely absent in the UHGG database.

To assess quantitative agreement between metagenomic and 16S data, we compared abundances of individual taxa. Across all samples, relative abundances at both the phylum and family levels were strongly correlated between sequencing modalities (**Figure 1F**), with no evidence of systematic bias at the phylum level (**Figure S1A**). At the family level, metagenomics tended to estimate higher abundances of Clostridiaceae and lower abundances of Ruminococcaceae compared to 16S (**Figure S1B**), although the combined abundance of these two families was similar between modalities (**Figure S1C**), likely reflecting differences in taxonomic reassignment within the UHGG reference^31^. Overall, shotgun metagenomics robustly recapitulated 16S-derived ecological dynamics in humanized mice, while enabling species-and strain-level resolution in subsequent analyses.

We next assessed concordance in conventional, mouse-adapted microbiotas. In these samples, metagenomics captured a smaller fraction of the taxa detected by 16S and showed reduced agreement in relative abundance estimates (**Figure S1D–F**). These differences likely reflect use of the UHGG database, which was developed for human gut genomes. Due to our goal of comprehensively resolving strain-level responses, we focused subsequent analyses on humanized mice, for which reference-based metagenomic profiling performed robustly.

### Diverse patterns of species-level response to repeated ciprofloxacin treatments

To examine how antibiotic exposure reshapes gut communities at the species level, we first analyzed the humanized mouse experiment consisting of two five-day courses of ciprofloxacin separated by nine days of recovery (**Figure 1B**). Since mice within the same cage had similar community dynamics^14^, we aggregated metagenomic reads from mice within the same cage at each time point to improve the resolution of species quantification. Across three cages, we reliably detected 218 species in total and classified each species’ response to each ciprofloxacin treatment using quantitative criteria based on changes in relative abundance (**Methods**). During each of the two consecutive treatments, species were assigned to five response classes based on their relative abundance dynamics (**Figure 2A**): 1) persistently remained low abundance (“low abundance”), defined as <0.1%, for which responses could not be reliably interpreted, 2) exhibited a marked decrease upon treatment and did not recover to pre-treatment levels afterward (“disrupted and not recovered”), 3) transiently increased during treatment (“transiently increased”), 4) decreased substantially and then recovered to near pre-treatment levels (“disrupted and recovered”), and 5) remained stable throughout treatment (“maintained”) (**Methods**). We interpret maintenance or increases in relative abundance during treatment as indicative of relative resistance or drug tolerance, whereas decreases reflect relative susceptibility within the community context.

**Figure 2:**
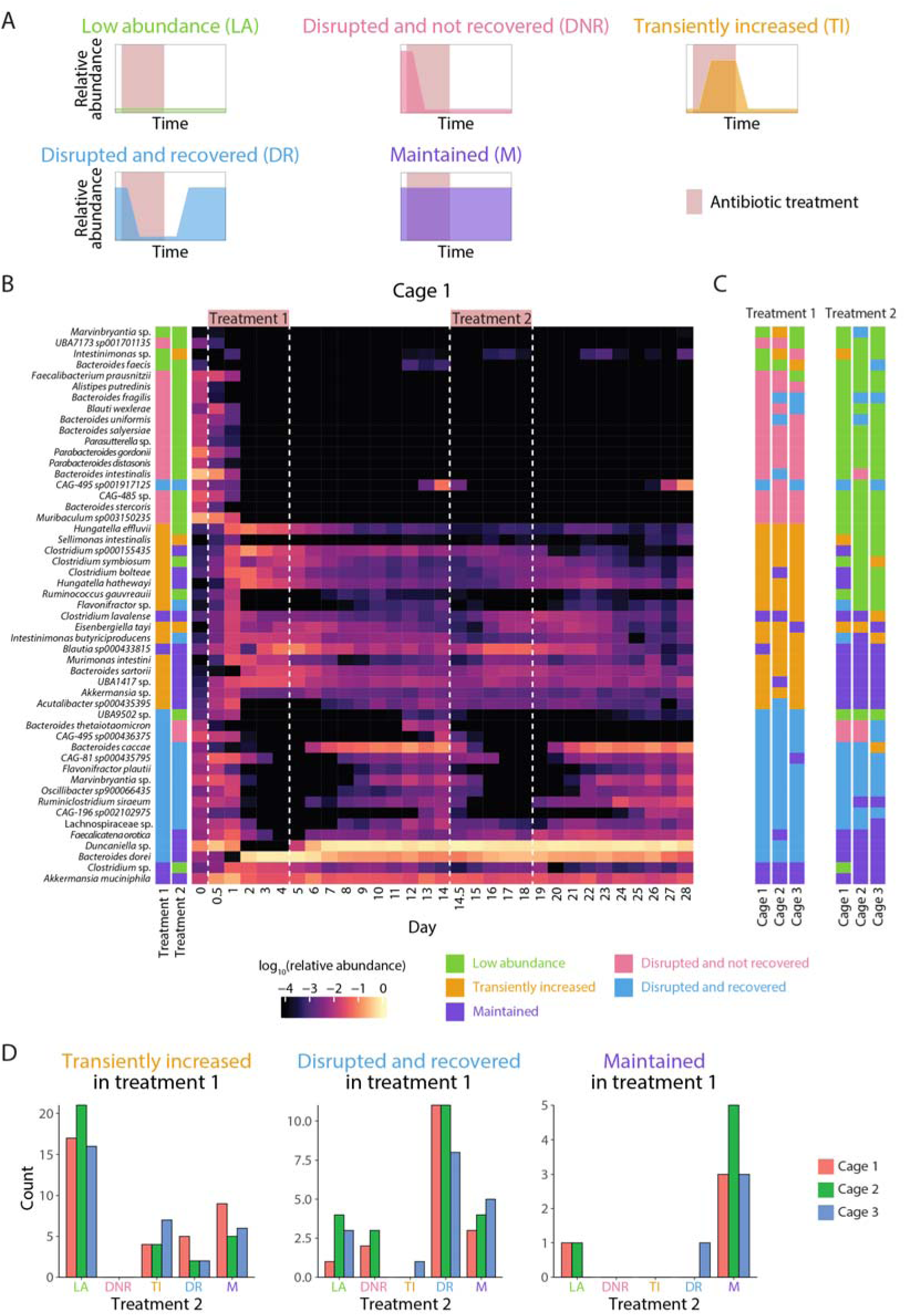
Diverse and path-dependent bacterial responses to repeated ciprofloxacin exposure. (A) Schematics for the five categories of response to an antibiotic treatment. (B) Heatmap of abundances for high-abundance species (maximum relative abundance > 1%) in cage 1 across the double ciprofloxacin treatment experiment. Species responses varied, with many remaining disrupted and a few showing resistance during the second treatment. Vertical dashed lines denote antibiotic treatment windows. (C) Heatmaps of species response classifications for the three cages stratified by treatment. Response patterns were largely consistent across cages in both treatments. Species ordering is matched between (B) and (C). (D) Bar plots of species response classification to the second ciprofloxacin treatment conditioned on the first treatment. Response to the second treatment can be qualitatively different from the first, potentially due to acquiring resistance mutations and/or changes in community context. Response to ciprofloxacin is relatively reproducible across cages. See also Figure S2, S3.

During the first ciprofloxacin treatment, most detected species were classified as low abundance (99/186 species in cage 1, 53%). Among the remaining higher-abundance species, there were similar numbers classified as disrupted and not recovered (31/186, 17%) or transiently increased (35/186, 19%), while fewer species were disrupted and recovered (17/186, 9%) and only a small number were maintained (4/186, 2%) (**Figure 2B**). Response patterns were broadly consistent across cages, and we present cage 1 as a representative example (**Figure 2C**). Across both treatments, the most common response patterns were low abundance in both treatments (95/186, 51%) and disrupted and not recovered followed by low abundance (31/186, 17%), reflecting the long-lasting disruptive effects of ciprofloxacin on the microbiome. A small number of species (3/186, 1.6%), including *Akkermansia muciniphila*, were classified as maintained in both treatments, likely reflecting intrinsic resistance among these initially abundant species. Overall, these response classifications were broadly consistent with those obtained from 16S profiling (**Figure S2**), while metagenomics identified additional species in several response categories.

Species that recovered after the first treatment displayed heterogeneous outcomes upon re-exposure. Of the 17 species classified as disrupted and recovered” in the first treatment, a large fraction (11/17, 65%) was again classified as disrupted and recovered in the second, suggesting partial tolerance that permits regrowth once antibiotic pressure is removed (**Figure 2D**). Two species transitioned from disrupted and recovered to disrupted and not recovered, suggesting reduced resilience across treatments. In contrast, three species, including *Bacteroides dorei*, recovered after the first treatment and then maintained their abundance during the second treatment (disrupted and recovered → maintained), suggesting resistance emergence. One additional disrupted and recovered species was under the detection threshold prior to the second treatment and was classified as low abundance.

Species that transiently increased during the first treatment likely benefited from reduced competition. Increases in relative abundance generally persisted at timepoints after the total biomass recovered (as measured by CFU counts^14^), indicating that these blooms were not simply an artifact of compositional normalization. Instead, we conclude that TI species likely expanded into ecological niches vacated by disrupted species, before declining during recovery as previously disrupted species regrew and displaced them (**Figure 2B, S3**). Notably, of the 35 transiently increased species in the first treatment, five (14%) were classified as disrupted and recovered in the second, indicating that species responses can shift across repeated antibiotic exposures. This shift suggests that transient blooms during the first treatment may not simply reflect intrinsic resistance but can emerge from changing ecological interactions as the community restructures. Thus, successive antibiotic exposures can probe how prior perturbation reshapes species responses.

Together, these results show that species within the same gut community exhibit diverse and sometimes shifting responses to repeated antibiotic exposure. While some responses are reproducible across successive treatments with the same antibiotic, others diverge, potentially because of prior ecological disruption and adaptive change, motivating strain-resolved analyses to distinguish resistance evolution from context-dependent ecological and evolutionary effects.

### Reproducible sweeps in *Bacteroides dorei* due to standing genetic variation in gyrA

To identify the genetic mechanisms underlying species-level responses to antibiotics, we analyzed single nucleotide variants (SNVs) in all detected species across mice and time points using our metagenomics data. We refer to each species–cage combination as a cage population. SNVs were detected in 98 species across 231 cage populations, with 24% of these cage populations meeting quality thresholds for SNV-calling (**Methods**). Among these 55 high-quality cage populations, 87% contained at least one SNV sweep (defined as a SNV with an allele that shifted in frequency from <20% to >80% over time; **Methods**), indicating widespread strain-level genetic changes during antibiotic perturbation.

We first examined whether known resistance mutations could explain response dynamics. Mutations at position 83 of DNA gyrase subunit A (encoded by *gyrA*), typically a serine residue, are well established determinants of ciprofloxacin resistance^32^. *B. dorei* recovered rapidly from the first ciprofloxacin treatment in all cages, reaching high relative abundance by day 3, and remained abundant throughout the second treatment (**Figure 3B**). This recovery coincided with the sweep of the *gyrA^S83F^* allele, a mutation previously shown to confer strong resistance^16^. Nearly 10,000 nonsynonymous SNVs displayed allele-frequency trajectories tightly correlated with *gyrA^S83F^*, indicating that the sweeping genotype represents a pre-existing strain background present in the initial community at low abundance. The reproducible selection of the same *gyrA^S83F^*-associated genotype across all three independent cages (**Figure 3B**) strongly suggests that this resistance-associated allele was pre-existing before treatment rather than arising *de novo* during the treatment. We extended our analyses to another experiment in which humanized mice were fed a MAC-deficient diet for two weeks prior to ciprofloxacin treatment (**Figure 3A**). A similar sweep of the *gyrA^S83F^*-containing strain was observed, indicating that ciprofloxacin-driven selection of this variant is robust to dietary context (**Figure 3C**).

**Figure 3:**
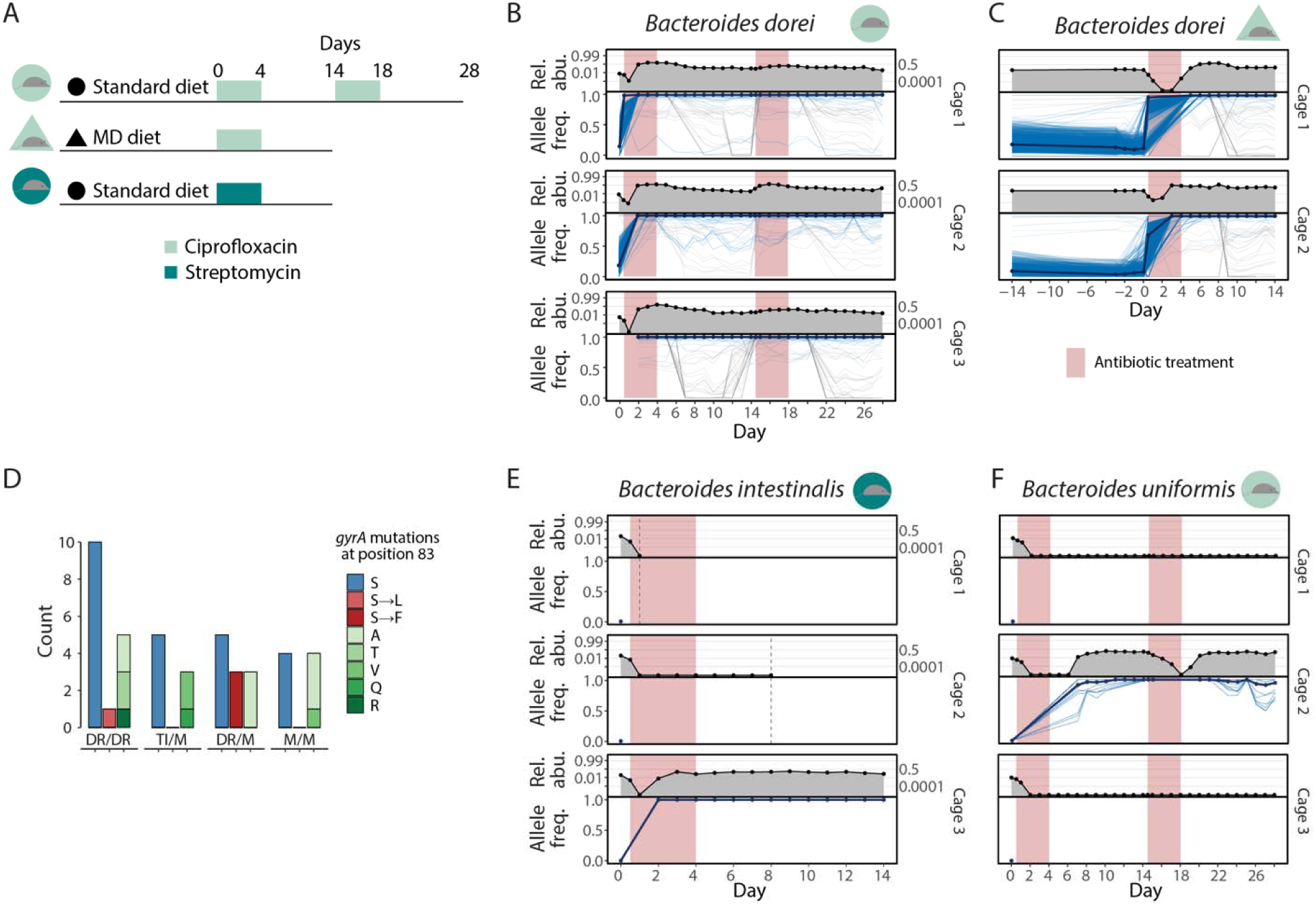
Antibiotic exposure can select for canonical resistance mutations. (A) Schematic of experimental conditions, including double ciprofloxacin treatment under a standard diet (SD), single ciprofloxacin treatment under a MAC-deficient diet (MD), and single streptomycin treatment under a SD. (B) Relative abundance (top) and SNV allele frequencies (bottom) for *B. dorei* over time in three cages in the double ciprofloxacin/SD experiment. Reads from mice within each cage were aggregated at each timepoint to improve detection of low-frequency variants. A pre-existing *B. dorei* strain carrying the canonical *gyrA^S83F^* mutation underwent a selective sweep in all cages. (C) In the single ciprofloxacin/MD diet experiment, *B. dorei* exhibited similar sweeps of pre-existing *gyrA* variants in both cages, indicating that this resistance mechanism is robust to dietary perturbation. (D) Distribution of DNA gyrase A alleles at position 83 across species, grouped by response category. Only four mutational sweeps were observed (one S83L and three S83F), while in most cases alleles did not exhibit selective sweeps. Notably, many species showing apparent resistance dynamics (TI/M and M/M) retained the wild-type serine allele, indicating that non-canonical mechanisms frequently underlie ciprofloxacin resistance. (E) *B. intestinalis* in the streptomycin/SD experiment exhibited a a *de novo* mutation in ribosomal protein S12 (K43R) in cage 3, consistent with canonical streptomycin resistance. (F) *B. uniformis* in the double ciprofloxacin/SD experiment carried a pre-existing *gyrA^S83L^*mutation (cage 2), yet remained susceptible to the second treatment, demonstrating that canonical resistance mutations are not always sufficient for persistence. In (B,C,E,F), pink shaded regions denote antibiotic treatment periods. Individual SNV trajectories are colored by clusters defined by temporal dynamics, and thick lines indicate the median trajectory of clusters containing canonical resistance mutations. Vertical dashed lines indicate time points at which the mice were sacrificed.

Despite this clear example, canonical *gyrA* mutations accounted for only a minority of resistance-associated recoveries. Among 11 species occurrences (by cage) classified as acquiring resistance (disrupted and recovered → maintained), only three (27%) developed *gyrA^S^*^83^ mutations (**Figure 3D**). Moreover, among species that either transiently increased or were maintained during the first treatment and maintained during the second, the wild-type serine allele remained the major allele throughout treatment in 62.5% (5/8) of transiently increased → maintained cases and 50% (4/8) of maintained → maintained cases. Thus, although selection on canonical *gyrA* resistance variants can drive rapid and reproducible recovery of *B. dorei* under ciprofloxacin treatment, these mutations only explain a subset of resistance-like responses.

#### *De novo* target-site mutation in ribosomal protein S12 drives streptomycin resistance in *Bacteroides intestinalis*

In mice humanized with the same donor fecal samples but treated with streptomycin (**Figure 3A**), which inhibits translation by binding the 30S ribosomal subunit, we observed multiple species in which sweeping strains carried canonical resistance mutations in the gene encoding ribosomal protein S12. *Bacteroides intestinalis* provided a clear example: its post-treatment recovery was tightly correlated with the sweep of a K42R substitution in ribosomal protein S12, a well-characterized determinant of streptomycin resistance^33^. The absence of additional sweeping SNVs in this strain, together with its occurrence in only one of three cages subjected to streptomycin perturbation (**Figure 3E**), strongly suggests that this mutation arose *de novo* rather than through selection on standing genetic variation. The mutation initially arose in a single mouse and subsequently spread to the remaining four mice (**Figure 3E**), consistent with a single acquisition event followed by transmission.

More broadly, across the 19 species with sufficient coverage that exhibited resistance-like recovery during the streptomycin treatment, all presented sweeping SNVs in ribosomal protein S12, with 14 (74%) of these species harboring mutations that appeared to arise independently without accompanying mutations (**Methods**). Thus, under streptomycin treatment, resistance-like recovery was frequently associated with canonical mutations in ribosomal protein S12. Taken together, the ciprofloxacin and streptomycin experiments show that target-site resistance mutations are one common route to recovery under antibiotic perturbation in this system, although the evolutionary mode differs across cases and sometimes implicates additional genetic and/or ecological mechanisms.

### Antibiotic-driven genetic selection beyond canonical target-site resistance in Bacteroides uniformis and A. muciniphila

While canonical target-site mutations explained some resistance-associated recovery dynamics, not all such mutations were associated with clear resistance phenotypes. For example, in ciprofloxacin-treated mice, *Bacteroides uniformis* exhibited cage-dependent dynamics: following disruption during the first treatment, no recovery was observed in cages 1 and 3, whereas in cage 2 *B. uniformis* rebounded to abundances exceeding pre-treatment levels. In cage 2, a *gyrA^S83L^* allele, previously shown to confer resistance in *Escherichia coli*^34^, fixed in the population by day 7 (**Figure 3F**). The presence of numerous tightly linked passenger alleles indicates selection on a pre-existing strain.

However, despite fixation of the S83L allele, *B. uniformis* remained susceptible during the second ciprofloxacin treatment, demonstrating that this mutation was not sufficient to enable robust persistence during the second ciprofloxacin treatment.

While most species exhibited similar dynamics across mice in the same cage, *A. muciniphila* displayed substantial heterogeneity across co-housed mice. *A. muciniphila*, a prevalent gut commensal in humans, has been reported to exhibit ciprofloxacin resistance across many independent isolates^35–37^. While *A. muciniphila* remained resistant across both treatments in a subset of mice in every cage, susceptibility varied within cages. In each cage, at least one mouse showed a decline in *A. muciniphila* abundance from >30% to below 1% during the first treatment (**Figure 4A**), and in three mice (two in cage 1 and one in cage 3), it fell below the limit of detection, indicating that *A. muciniphila* exhibits variation in observed resistance across hosts.

**Figure 4:**
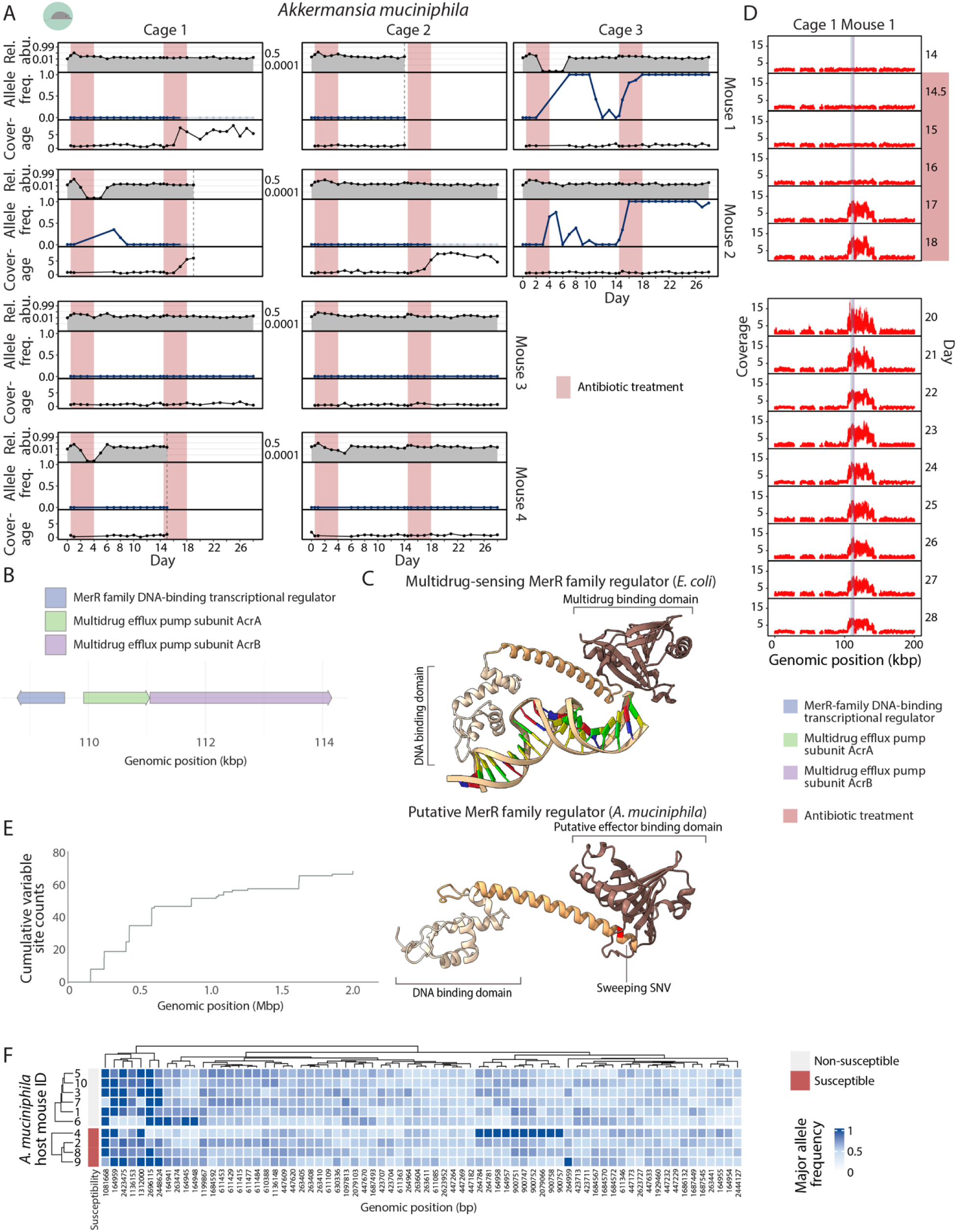
Distinct genetic mechanisms underlie heterogeneous ciprofloxacin susceptibility in *Akkermansia muciniphila*. (A) Relative abundance (top), allele frequencies of SNVs (middle), and relative coverage normalized by the median coverage across all genomic sites (bottom) of *A. muciniphila* over time in individual mice across three cages. A single SNV sweeping in both mice from cage 3 maps to a MerR-family transcriptional regulator, with both allele fixation and local amplification tightly associated with ciprofloxacin treatment. However, resistance in other mice lacking this mutation indicates the presence of alternative genetic mechanisms. Shaded areas denote ciprofloxacin treatment periods, and vertical dashed lines denote time points at which mice were sacrificed. Allele frequencies are shown for all time points, with time points exceeding a relative coverage threshold of 3 displayed with increased transparency. (B) Genomic organization of the MerR transcriptional regulator and its putative targets in the reference *A. muciniphila* genome (RefSeq assembly GCF_902387255.1), including the AcrA and AcrB multidrug efflux pump subunits. (C) Structural comparison of the multidrug-sensing MerR-family regulator from *E. coli* (top) and the putative *A. muciniphila* homolog predicted using AlphaFold2 and HMMER (bottom). DNA-binding and effector/drug-binding domains are annotated. Conserved structural features, including the multidrug-binding domain, support a potential role for this regulator in antibiotic response. (D) Relative coverage profiles across the *A. muciniphila* genome (0–200 kbp) in a mouse during the second ciprofloxacin treatment. Points denote individual genomic positions, highlighting putative amplification events. This mouse is one of three individuals that exhibit amplification of this region, which contains three genes tightly linked to ciprofloxacin resistance. Amplification emerged around day 17, during the second treatment, and persisted through the remainder of the experiment. Co-amplification of the MerR regulator and adjacent efflux pump genes suggests a coordinated genetic mechanism that may contribute to reduced antibiotic susceptibility. Time points lacking data correspond to low-abundance samples with insufficient coverage for reliable SNV detection. (E) Cumulative distribution of 77 variable sites identified in the seeding population of *A. muciniphila* across all mice. (F) Heatmap of consensus allele frequencies at variable sites across *A. muciniphila* strains isolated from mice. Variable sites were hierarchically clustered by allele distribution across strains, and strains were clustered by variant profiles within susceptibility groups. No clear association was observed between genotype and apparent ciprofloxacin susceptibility phenotype. See also Figure S4.

In both of the two mice from cage 3, a single allele in *A. muciniphila* exhibited a striking trajectory tightly coupled to antibiotic exposure: it rose from undetectable levels during the first treatment, declined back to undetectable levels during recovery, and then fixed at 100% during the second treatment (**Figure 4A**). This allele maps to residue 118 of a putative MerR-family DNA-binding transcriptional regulator. The regulator is adjacent to the multidrug efflux pump subunits *acrA* and *acrB*, strongly implicating a role in antibiotic response (**Figure 4B**). Domain analysis using HMMER (**Methods**) revealed an N-terminal helix-turn-helix DNA-binding domain, and structural modeling with AlphaFold2 showed close similarity to the *E. coli* MerR-family multidrug-sensing transcriptional regulator determined by cryo-electron microscopy^38^ (**Figure 4C**). The sweeping allele localizes to the linker region connecting the N-terminal DNA-binding and C-terminal effector-binding domains, suggesting altered regulatory function. Notably, in one mouse in which this allele swept, the *A. muciniphila* population appeared ciprofloxacin resistant prior to treatment, indicating that antibiotic exposure can drive rapid strain-level turnover even without substantial population collapse. Conversely, *A. muciniphila* populations in some mice in other cages remained resistant across both treatments despite lacking this allele.

In addition to the regulatory mutation, we observed evidence for a distinct genetic route to ciprofloxacin resistance involving local genomic amplification in *A. muciniphila*.

Elevated relative coverage ratio (**Methods**) spanning the MerR-family regulator and adjacent *acrA*/*acrB* efflux pump genes was detected in three mice, two from cage 1 and one from cage 2 (**Figure 4D, S4**). In each of these mice, the amplification emerged midway through the second treatment and persisted thereafter. The coordinated timing and reproducibility of this amplification across mice suggest that increased copy number of the regulator–efflux pump module provides an alternative mechanism of resistance.

These amplifications were detected only during the second ciprofloxacin treatment, highlighting how prior perturbation history can open additional evolutionary routes upon re-exposure. However, the MerR regulator did not account for all resistant *A. muciniphila* populations, as *A. muciniphila* in other mice appeared ciprofloxacin resistant despite lacking both the MerR-associated mutation sweep and amplification.

Together, these findings demonstrate that antibiotic exposure can select a diverse range of genetic changes, including mutations that do not directly confer resistance, mutations whose effects are context dependent, and multiple alternative resistance mechanisms within a single species. Such complexity highlights that genetic sweeps observed during antibiotic treatment cannot be interpreted solely through the lens of canonical resistance mutations, but instead reflect a broader landscape shaped by ecological context and strain-level variation.

### Heterogeneous susceptibility of *A. muciniphila* cannot be explained by SNV differences between the seeding populations

Across replicate mice exposed to the same perturbation, *A. muciniphila* showed heterogeneous strain-level responses: among the eight mice with reduced susceptibility to the second ciprofloxacin treatment, three lacked evidence of strain-level sweeps or local genomic amplification (cage 1, mouse 3; cage 2, mouse 3; cage 2, mouse 4). We therefore asked whether the observed heterogeneity in ciprofloxacin susceptibility could be attributed to SNV differences that accumulated between the initial seeding populations.

To evaluate this possibility, we compared the major allele at all genomic sites with >10X coverage in each mouse at 12 h post-gavage with ciprofloxacin. We selected this time point rather than baseline (*t*=0) because the higher relative abundance of *A. muciniphila* at 12 h provided improved per-site coverage for cross-mouse comparisons. Across the ∼2.3 Mbp *A. muciniphila* genome, we identified only 77 variable sites among the 10 mice, corresponding to >99.99% sequence identity among dominant strains (**Figure 4E**). With a single triallelic exception (site 244127 in mouse 9), all sites were monoallelic or biallelic, and in the latter case the pair of alleles was observed across mice. We defined the allele present as the majority in more than five mice as the consensus allele and quantified its frequency at each variable site (**Figure 4F**). At many sites, the consensus and alternate alleles were present at similar frequencies across mice, indicating that a shared mixture of alleles underlies these populations.

Despite this shared genetic background, allele identities at these sites showed no association with susceptibility phenotype, indicating that variation in ciprofloxacin response cannot be explained by SNVs in the dominant strains of the seeding population (**Figure 4F**). Although this analysis rules out a primary role for single nucleotide differences, it does not exclude contributions from other forms of genetic variation, including inversions, insertions or deletions, plasmid carriage, or copy number variation. Together, these results suggest that antibiotic susceptibility of gut commensals *in vivo* is not necessarily predictable from SNVs alone, highlighting the complexity of drug–microbiome interactions.

### Parallel antibiotic exposure yields distinct fixation events in *Duncaniella* species across cages and individual mice

Another example of strain-level responses that varied across and within cages was observed in a *Duncaniella* species, which exhibited striking cage-specific and mouse-specific fixation events under parallel ciprofloxacin exposure. Across all mice and cages, *Duncaniella* populations were substantially less susceptible during the second treatment than during the first, consistent with the emergence of resistance. However, the underlying genetic changes associated with this phenotype differed markedly across cages and even among mice within the same cage.

In cage 1, all four mice exhibited a sweep of a nonsynonymous SNV in the gene encoding the efflux-pump membrane transporter BepE (**Figure 5A**). This sweep was absent in the other cages, indicating that the *bepE* mutation is not required for the reduced susceptibility observed during the second treatment. In contrast, in cage 3, we observed sweeps of nonsynonymous variants mapping to genes without an obvious connection to antibiotic resistance, including asparagine synthetase B (*asnB*) and the sensory transduction regulator *lytR* (**Figure 5A**). These cage-specific fixation events suggest that similar resistance-associated phenotypes can arise through distinct and potentially indirect genetic trajectories.

**Figure 5:**
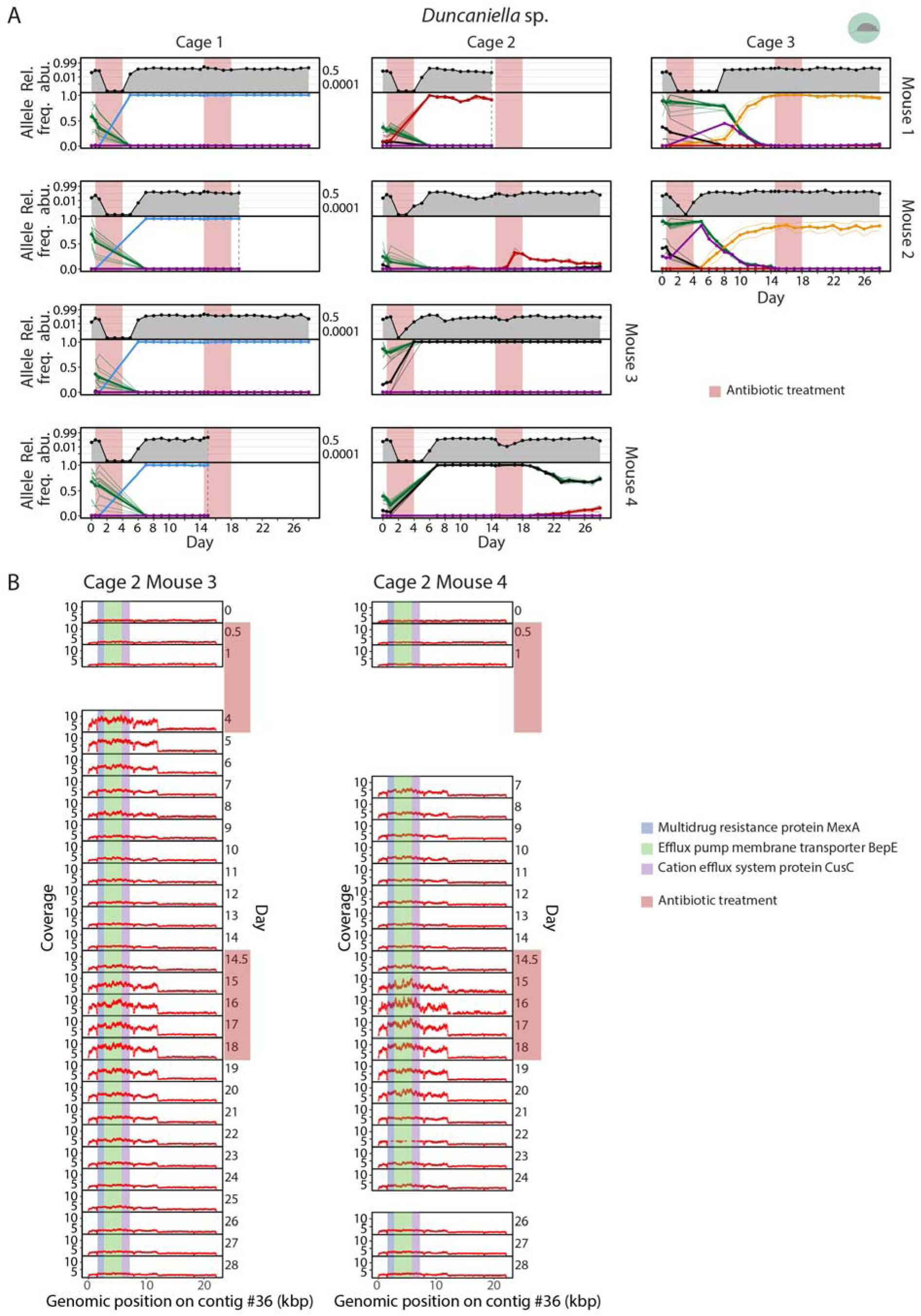
Parallel ciprofloxacin exposure drives divergent strain-level fixation events in *Duncaniella*. (A) Relative abundance (top panels) and allele frequencies of SNVs (bottom panels) are shown for *Duncaniella* species over time in individual mice across three cages. Responses to antibiotics varied across mice, including within the same cage, highlighting the stochastic nature of strain-level dynamics. SNV traces are colored according to clusters defined by their temporal dynamics. The six clusters each contain nonsynonymous SNVs from the following genes. Blue: efflux pump membrane transporter BepE and adaptive-response sensory-kinase SasA; red: efflux pump membrane transporter BepE and amidophosphoribosyltransferase; green: DNA ligase, sensor histidine kinase RcsC, TonB-dependent receptor P3, TonB-dependent receptor SusC, ribosomal RNA small subunit methyltransferase F, and starch-binding protein SusD; black: TonB-dependent receptor P3 and fumarate hydratase class I; orange: asparagine synthetase B and sensory transduction protein LytR; and purple: TonB-dependent receptor P26. Selective sweeps of pre-existing variants occurred in genes associated with antibiotic resistance as well as in genes with no obvious connection to resistance. Shaded regions indicate ciprofloxacin treatment periods, and vertical dashed lines denote time points at which mice were sacrificed. (B) Relative coverage profiles along contig 36 of the *Duncaniella* sp. genome in two mice across both ciprofloxacin treatments. Points denote individual genomic positions, highlighting putative amplification events. Colored regions indicate loci encoding efflux-associated proteins, including MexA, BepE, and CusC. Shaded regions denote antibiotic treatment periods. These two mice are the only individuals exhibiting amplification of this region. Amplification emerged during the first treatment, persisted transiently, and reappeared rapidly at the start of the second treatment, consistent with treatment-coupled, repeatable amplification dynamics. These amplification events were not present in other mice within the same cage, indicating substantial stochasticity at the strain level. Time points lacking data correspond to low-abundance samples with insufficient coverage.

In addition to single nucleotide changes, we observed treatment-coupled genomic amplification in *Duncaniella*, analogous to the amplification events observed in *A. muciniphila* (**Figure 5B**). In cage 2, only two of the four mice (mice 3 and 4) exhibited increased copy number across a focal genomic region containing the efflux-associated genes *bepE*, *mexA*, and *cusC*. This amplification emerged midway through the first ciprofloxacin treatment, disappeared during recovery, and re-emerged immediately upon initiation of the second treatment, persisting throughout antibiotic exposure before returning to baseline afterward. The reproducibility and tight coupling of this amplification to antibiotic exposure suggest that increased gene copy number of efflux-related genes provides an alternative resistance mechanism in *Duncaniella*.

Notably, strain-level responses also varied substantially among mice within the same cage. In cage 2, some SNVs swept during the first treatment in mice 3 and 4 but not in mice 1 and 2, whereas other SNVs increased in frequency during the first treatment only in mouse 1 and remained approximately constant in the remaining mice (**Figure 5A**). Moreover, only mice 3 and 4 exhibited the efflux-associated amplification, further illustrating the heterogeneity of evolutionary trajectories even among co-housed animals and suggesting that within-cage transmission or establishment of this amplified lineage was limited.

Together, these observations demonstrate that parallel antibiotic exposure can yield divergent strain-level evolutionary outcomes across cages and even among individual mice. Such variability underscores the stochastic nature of strain-level dynamics in the gut microbiome. Although dense longitudinal metagenomics can resolve specific candidate mechanisms, including SNV sweeps and genomic amplifications, these examples likely capture only a subset of the genetic routes underlying the reduced antibiotic susceptibility observed in this system.

### Strain transfer between two humanized mouse cohorts during and after antibiotic treatment

In addition to ecological and evolutionary changes of resident species, invasion of non-resident strains can shape antibiotic response. To investigate strain transfer between hosts during antibiotic treatment, we analyzed an experiment in which mice were humanized with stool from two distinct donors (H1 and H2). H1 corresponds to the donor microbiota used in the previous humanized mouse experiments. After microbiota equilibration, a subset of mice was cross-housed at defined times relative to the first ciprofloxacin treatment: immediately before treatment (day 0), during treatment (day 1), or immediately after treatment (day 5). As controls, some H1- and H2-humanized mice remained separately co-housed throughout the experiment (**Figure 6A**). To distinguish donor-derived strains following cross-housing, we identified cohort-specific SNVs for each species shared between H1 and H2 mice and used these SNVs as markers to assign strain origin (**Methods**). Species-level changes after cross-housing may reflect true donor-strain transfer, recovery of low-abundance resident strains, or both, necessitating strain-resolved analysis to distinguish among these possibilities.

**Figure 6:**
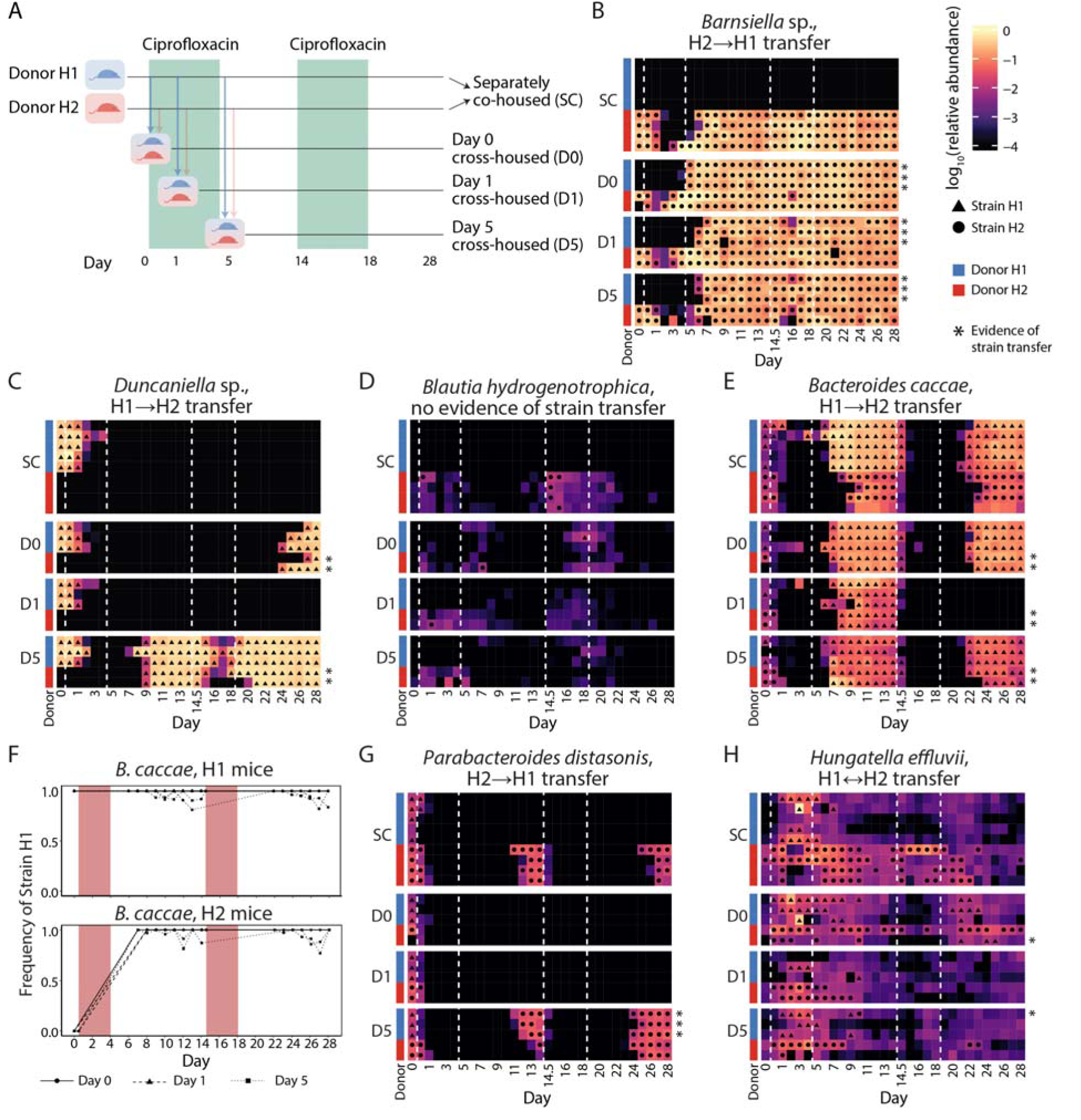
Strain-resolved metagenomics reveals bidirectional strain transfer and replacement during post-antibiotic recovery. (A) Schematic of the cross-housing experiment. Mice harboring humanized microbiota from two donors (H1 and H2) were either separately co-housed with mice from the same human donor (SC) or cross-housed with mice from the other donor at defined times relative to antibiotic treatment (day 0, 1, or 5; D0, D1, D5). (B) Heatmap of *Barnsiella* species across mice and time points, showing transfer of the H2-originating strain into H1 mice across all cross-housing groups. (C) For a *Duncaniella* species, the H1-originating strain transferred into H2 mice in the D0 and D5 cross-housing groups.(D) For *Blautia hydrogenotrophica*, the strain detected in H1 mice after cross-housing is distinct from the H2-originating strain (e.g., in the D0 group), suggesting recovery of a resident strain rather than true transfer and highlighting the importance of strain-resolved analysis. (E) The H1-originating *B. caccae* strain replaced the resident strain in H2 mice following the first treatment across all cross-housing groups. The transferred strain was maintained after the second treatment in the D0 and D5 groups while *B. caccae* did not recover in the D1 group. (F) Frequency of the H1 strain in *B. caccae* is shown over time in both H1 and H2 mice. In H2 mice, the incoming H1-derived strain fully displaced the resident strain across all three cross-housing cohorts, whereas in H1 mice, the H1-derived strain was consistently maintained throughout the experiment. Each trajectory represents an individual mouse, labeled by D0, D1, or D5 cross-housing group. (G) The H2-originating *P. distasonis* strain replaced the resident strain in H1 mice in the D5 cross-housed group and was maintained after the second treatment. (H) In a single case (*Hungatella effluvii*), strain replacement occurred without prior loss of the resident strain, observed in one H1 mouse in the D5 cross-housed group. In (B-E, G-H), each mouse is annotated by its initial donor (H1 or H2). Strain identities (black circles and triangles) are shown for samples with sufficient coverage for SNV calling. Strain assignment was determined using donor-specific marker SNVs identified from separately co-housed mice (**Methods**), with H1 strains characterized by SNPs conserved across SC H1 mice but absent in SC H2 mice, and vice versa. The exception was for *B. hydrogenotrophica* in (D), where the H1 strain was identified only in one of the H1 mice in the D0 cross-housed group. Vertical dashed lines denote antibiotic treatment windows.

Prior 16S profiling of this cross-housing experiment suggested that a *Barnesiella* species transferred from H2 to H1 mice: the ASV representing this species was abundant in H2 mice but undetectable in separately co-housed H1 mice, and it appeared at high abundance in H1 mice after cross-housing^14^. However, such species-level data could not conclusively establish whether this observation reflected transfer of a strain in the H2 cohort rather than the emergence of a strain in the H1 cohort from below the limit of detection. Our strain-resolved metagenomics analysis showed that the *Barnesiella* population appearing in H1 mice did in fact carry H2-specific SNV markers, confirming transfer of the H2 strain (**Figure 6B**).

A complementary pattern was observed for the previously discussed *Duncaniella* species (**Figure 5**), which was present in separately co-housed H1 but absent from H2 mice. In cohorts cross-housed on day 0 or day 5, the H1-derived *Duncaniella* strain emerged in H2 mice (**Figure 6C**). Donor-specific SNVs confirmed that the transferred strain originated from the H1 cohort. The timing of its appearance differed between cohorts. In mice cross-housed on day 0, the H1-derived *Duncaniella* strain appeared in H2 mice only after the second antibiotic treatment, whereas in mice cross-housed on day 5 it appeared after the first treatment. In contrast, no transfer was observed in mice cross-housed on day 1. These differences mirror the recovery dynamics of *Duncaniella* in H1 mice: recovery was delayed in the day 0 cohort and absent in the day 1 cohort, suggesting that strain transfer depends on donor population recovery following antibiotic exposure. These results are consistent with the stochastic recovery dynamics described earlier for this *Duncaniella* species (**Figure 5**).

In contrast to these clear transfer events, apparent transfer at the species level did not always reflect transfer of the dominant donor strain. In the control mice that were never cross-housed, *Blautia hydrogenotrophica* was detected only in H2 mice. After cross-housing, *B. hydrogenotrophica* was detected in many H1 mice, albeit at low abundance and with substantial inter-mouse variability (**Figure 6D**). However, strain-resolved analysis revealed that the *B. hydrogenotrophica* population in H1 mice differed from the H2 strain at 395 SNV sites distributed broadly across the genome. This divergence indicates that the detected *B. hydrogenotrophica* population reflects either expansion of a low-abundance resident strain in H1 mice or transfer of a distinct, minor strain from H2 mice. In either case, strain-resolved metagenomics was essential for distinguishing true donor-strain transfer from apparent species-level transfer.

### Strain-level analysis distinguishes recovery of resident strains from cross-cohort strain transfer

For species that were disrupted but later appeared to recover in the cross-housing experiment, 16S rRNA sequencing alone cannot determine whether post-treatment recovery reflects regrowth of the original dominant strain, expansion of a different resident strain, or acquisition of a strain transferred between mice – particularly when the species is present in both H1 and H2 cohorts. *Bacteroides caccae* illustrates this ambiguity. This species was detected in all H1 and H2 mice prior to the first ciprofloxacin treatment, decreased below the limit of detection during treatment, and subsequently recovered. However, across all three cross-housing cohorts, the dominant *B. caccae* strain in H2 mice was replaced by the H1-derived strain during recovery, as revealed by strain-specific SNV markers (**Figure 6E**). During the second antibiotic treatment, the H1 strain again declined below the limit of detection and subsequently recovered in all cohorts except mice cross-housed on day 1, in which *B. caccae* abundance remained below the limit of detection in all five mice in that cage. When this strain replacement occurred, the incoming H1-derived strain fully displaced the initial strain in H2 mice across all three cross-housing cohorts (**Figure 6F**).

A similar pattern was observed for other species, although strain replacement was sometimes cage dependent. In the control mice, *Parabacteroides distasonis* recovered after the first antibiotic treatment in H2 mice but remained below the level of detection in H1 mice, suggesting a trajectory toward extinction in the latter cohort. Among cross-housed cohorts, *P. distasonis* recovered only in mice cross-housed on day 5. In these animals, the H2-derived strain recovered in H2 mice after the first treatment and was also detected in H1 mice (**Figure 6G**), consistent with strain transfer. This strain again dropped below the limit of detection during the second treatment and then recovered thereafter.

Together, our strain-resolved metagenomic analyses show that apparent species-level recovery can reflect either regrowth of resident strains or engraftment of transferred strains, indicating that post-antibiotic transmission is conditional and often drives competitive strain replacement rather than restoration of the original population.

### Strain replacement can occur without antibiotic-induced elimination of the resident strain

In *Hungatella effluvii*, our replicated experimental design revealed variability in strain transfer outcomes even across mice within the same cage. In the day 0 cross-housing cohort, the resident strain in one H2 mouse declined below the limit of detection during the second ciprofloxacin treatment and was subsequently replaced by the H1-derived strain. In contrast, in a cage mate exposed to the same conditions, the H2 strain never decreased below the limit of detection during either treatment, and no strain replacement was observed (**Figure 6H**).

Although strain replacement was typically preceded by apparent elimination of the resident strain, we identified a rare instance in which replacement occurred without prior loss of the resident population. In one H1 mouse in the day 5 cross-housing cohort, *H. effluvii* increased in relative abundance during the first treatment and retained the H1-specific marker SNVs through day 9. However, on day 10, the H1 strain was abruptly and completely replaced by the H2 strain (**Figure 6H**). This observation demonstrates that strain replacement can occur rapidly and independently of antibiotic-induced extinction of the resident strain.

Overall, these results indicate that strain transfer between H1 and H2 mice is bidirectional and is frequently, but not invariably, associated with prior loss of the resident strain. In all strain replacement examples discussed here, the incoming strain effectively displaced the resident strain, with no evidence of stable coexistence over the time scales examined (**Figure 6F**).

### Phage activity is heterogeneous across mice but tends to cluster within cage

Bacterial genomes often harbor prophage elements that respond to antibiotic challenge through induction. At the same time, many prophages in the human gut are thought to persist primarily in a lysogenic state, contributing little detectable free-phage activity under steady-state conditions^39,40^. To determine whether phage dynamics track bacterial population changes during the antibiotic perturbation, we analyzed longitudinal viral abundance profiles alongside bacterial abundance using Phanta^39^ (**Methods**), a *k*-mer-based classifier that estimates bacterial and viral genome abundances from short-read data. We focused on the two-course ciprofloxacin experiment in humanized mice (**Figure 1B**). Rarefaction analysis of day 0 samples revealed a steeper rarefaction curve for phages than for bacteria (**Figure S5A**), consistent with smaller phage genome sizes and higher apparent diversity for a given sequencing depth. Across all samples, the genomic virus-to-microbe ratio (gVMR) was 3.6±2.0 (mean±S.D.; **Figure S5B**), comparable to values reported in human studies^39,40^, supporting the suitability of this humanized mouse system for interrogating bacteria-phage dynamics in the gut microbiome.

To characterize temporal relationships between phages and bacteria, we categorized phage trajectories into four classes based on cross-correlation with bacteria abundance profiles. First, some phages exhibited little temporal structure (group I, **Figure 7Ai**), consistent with low-level activity or hosts below the detection threshold. Second, a subset of phages closely mirrored the abundance of a specific bacterial species (group II, **Figure 7Aii**), suggesting tight coupling consistent with lysogenic inheritance. Third, some phages displayed dynamic changes with weak or no correlation to any single bacterial trajectory (group III, **Figure 7Aiii**), indicating activity not simply explained by host abundance (potentially lytic). Finally, a fourth class partially tracked the abundance of a bacterial species but exhibited additional spikes that deviated from bacterial dynamics (group IV, **Figure 7Aiv**), suggestive of episodic activity beyond passive lysogenic replication. For groups II and IV, putative bacterial hosts were assigned by identifying the strongest correlation between bacterial and phage abundances across mice and time points (**Methods**). Across the 87 phages analyzed, five were classified as group III and 14 as group IV, together comprising ∼22% of phages exhibiting measurable dynamic activity (**Figure 7B**).

**Figure 7:**
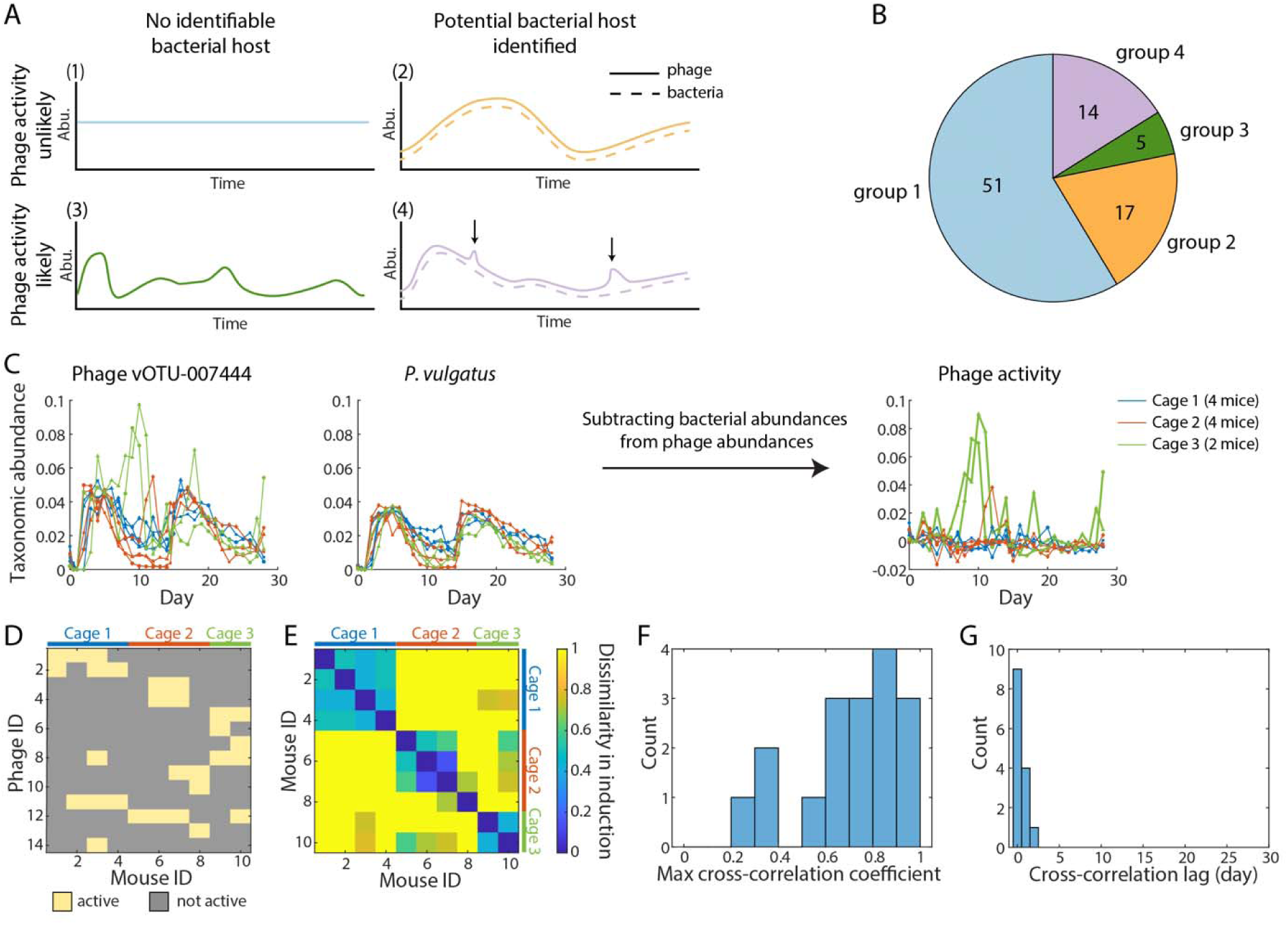
Phage activity during antibiotic perturbation is heterogeneous and clusters within cages. (A) Schematic of phage group classification based on temporal dynamics and correlation with potential bacterial hosts. (B) Distribution of phages across groups identified in the double ciprofloxacin treatment experiment. (C) Representative example of phage vOTU-007444, whose abundance tracks *P. vulgatus* in cages 1 and 2 but exhibits additional activity in cage 3 that is not explained by host abundance, consistent with induction of other lytic processes. (D) Heatmap of phage activity events for all group IV phages across mice. Activity is heterogeneous across mice but shows clustering within cages. (E) Dissimilarity matrix of phage activity profiles across mice, quantified using Bray-Curtis distance. Mice within the same cage exhibit more similar activity profiles but typically diverge across cages. (F) Maximum cross-correlation coefficients of phage activity among cage mates sharing active phages. In 14/17 pairs, phage activity was correlated (Pearson’s *r*>0.5). (G) Cross-correlation lags for the 14 pairs with *r*>0.5. Lags are near zero, indicating synchronized phage activity within cages. See also Figure S6.

Given the distinct dynamics of the phage groups, we quantified and compared the temporal variability of phages in groups II, III, and IV^41^. All three groups exhibited power-law scaling between the mean and variance of phage abundances, with group III showing the largest exponent, and group IV an intermediate exponent (**Figure S5C**), consistent with their more pronounced and irregular fluctuations over time. We next assessed inter-mouse heterogeneity by calculating the coefficient of variation (CV) across mice at each time point for both phages and bacteria (**Figure S5D**). Group II phages exhibited CVs comparable to their bacterial hosts (*p*=0.96, one-tailed Kolmogorov-Smirnov test), consistent with tight host coupling through lysogeny. In contrast, group IV phages showed significantly greater heterogeneity (*p*=0.001, one-tailed Kolmogorov-Smirnov test), and group III phages exhibited markedly elevated CVs relative to their bacterial hosts (*p*<10^-5^, one-tailed Kolmogorov-Smirnov test), indicating substantial mouse-specific dynamics. These patterns are consistent with the intermittent, host-dependent bursts of activity observed in groups III and IV, reflecting phage behaviors that extend beyond simple tracking of bacteria abundance.

We next focused on group IV phages to examine prophage-associated activity. A representative example is vOTU-007444, whose abundance closely tracked that of *Phocaeicola vulgatus* in cages 1 and 2 (**Figure 7C**). After rescaling and subtracting putative bacterial trajectory (**Methods**), residual phage abundance in these cages was minimal, consistent with largely host-linked dynamics. In contrast, cage 3 exhibited transient bursts of residual phage abundance that could not be explained by host fluctuations alone. Although these signals could in principle reflect activity on an alternative host, we did not observe any bacterial species with dynamics consistent with these residual peaks. Thus, these bursts are consistent with induction or other lytic processes.

Across all group IV phages, we quantified the ratio of phage to bacterial abundance during periods of activity and found that most events fell within a 1.3–3.6× range (25^th^–75^th^ percentile) relative to their putative hosts, consistent with prior long-read, assembly-based estimates of prophage induction in the human gut^42^. Together, these analyses demonstrate that longitudinal sampling combined with co-abundance modeling enables detection of low-level, transient phage activity in complex gut communities^40,42^.

Motivated by the variability of vOTU-007444 activity across cages, we further assessed phage activity in each mouse individually by constructing a binary matrix representing phage activity for group IV phages across mice (**Figure 7D**). A phage is considered active in a mouse host if it shows activity for at least one time point (**Methods**). Phage activity was highly heterogeneous across mice: although all mice were colonized with the same donor microbiota and underwent identical antibiotic treatment, individual phages were typically active in only one or a few mice (**Figure 7D, Figure S5E**).

Pairwise comparisons revealed that mice within the same cage shared more similar phage activity profiles than mice from different cages (**Figure 7E**). In 13/14 (93%) of group IV phages, activity was restricted to a subset of mice within a cage, while cage mates carrying the same phage exhibited no detectable activity. Among cage mates sharing active phages, residual phage trajectories exhibited high maximum cross-correlation coefficients (**Figure 7F**) with near-zero temporal lags (**Figure 7G**), indicating synchronized activity within cages.

We then quantified the temporal dynamics of prophage activity by counting the number of active prophages at each time point. Although ciprofloxacin has been reported to rapidly induce specific prophages in mice^43^, we did not observe a corresponding increase in activity among group IV phages. Instead, prophage activity exhibited a modest decrease during both ciprofloxacin treatments (**Figure S5F**), a pattern not explained by changes in bacterial host abundance (**Figure S5G**). These results suggest that the relationship between ciprofloxacin exposure and prophage induction is phage-specific and dependent on ecological context. Notably, our analysis captures only prophages detectable by Phanta within this experimental framework and therefore does not fully represent the diversity of inducible prophages in the gut microbiome. Improved methods using *de novo* phage genomes may reveal clearer interactions between antibiotic stress and phage induction. Together, these findings indicate that phage responses to antibiotic perturbation span a continuum from host-tracking lysogenic behavior to highly heterogeneous, cage-clustered bursts of activity, revealing an additional layer of variability to microbiome recovery.

## DISCUSSION

Dense longitudinal sampling of replicated, antibiotic-perturbed mouse microbiotas allowed us to dissect how closely matched communities exposed to the same perturbations can nonetheless follow divergent recovery trajectories. Across humanized mouse cohorts, some responses were reproducible across cages and mice, whereas others diverged markedly, indicating that post-antibiotic dynamics are more stochastic and path dependent than expected in this controlled setting. Species-level trajectories were consistently captured by both 16S rRNA profiling and metagenomic sequencing (**Figure 1**), while metagenomics further resolved the strain-level processes and phage dynamics underlying this divergence. We found that post-antibiotic recovery was shaped by selection on genetic variants (**Figure 3-5**), strain transmission between hosts (**Figure 6**), and activation of prophages (**Figure 7**). Together, these results show that antibiotics do not simply select for resistance within isolated populations, but instead initiate a cascade of interacting ecological, evolutionary, and mobilome processes that collectively govern microbiome reassembly.

At the genetic level, this cascade of processes manifests as a diverse set of strain-resolved evolutionary responses to antibiotic exposure that were not apparent from 16S profiling. Across species and treatments, we observed widespread selective sweeps during and after antibiotic perturbation, arising from both *de novo* mutation and selection on standing variation, some of which has previously been shown to confer resistance. In some cases, these sweeps involved canonical target-site resistance mutations, such as reproducible fixation of the *gyrA^S83F^* allele in *B. dorei* during ciprofloxacin treatment (**Figure 3B,C**) or *de novo* mutations in ribosomal protein S12 under streptomycin exposure (**Figure 3E**). These events closely parallel resistance dynamics described in humans^16^ and demonstrate that strong antibiotic pressure can rapidly drive fixation of high-effect variants even within complex gut communities. Consistent with observations in human cohorts^16^, resistance-associated variants in our system swept within days of antibiotic exposure and remained fixed well after treatment cessation, indicating that even brief perturbations can durably reshape the genetic landscape of gut commensals. Although our experiments were not designed to directly measure the fitness costs of these variants, their persistence after treatment suggests that some resistance-associated changes impose limited fitness costs *in vivo*. However, such classic resistance mutations accounted for only a subset of resistance-associated recovery events. Many species that recovered or persisted across treatments lacked known target-site mutations, indicating that reduced susceptibility frequently emerged through alternative genetic routes. We further observed local genomic amplifications and sweeps of mutations whose phenotypic effects were not canonical genetic determinants of resistance, highlighting that antibiotic-driven selection *in vivo* often acts on multiple genetic mechanisms simultaneously. For example, in *A. muciniphila*, we detected antibiotic-linked sweeps in a MerR-family regulator adjacent to the AcrA/AcrB efflux operon (**Figure 4A-C**), while in other mice increased copy number of the same operon was observed in the absence of this mutation (**Figure S4**), suggesting parallel but genetically distinct routes to reduced antibiotic susceptibility. Together, these findings show that while antibiotics can impose strong and repeatable selective pressures, the resulting genetic responses are mechanistically heterogeneous and extend beyond simple models of single-mutation resistance.

While genetic variation provides the raw material for selection, our results show that the evolutionary consequences of antibiotic exposure are strongly shaped by ecological context, rendering recovery trajectories path dependent. We observed multiple instances in which mutations swept during antibiotic treatment without conferring resistance in subsequent exposures, indicating that their selective advantage may depend on transient community configurations rather than drug tolerance alone. For example, in *B. uniformis*, fixation of *gyrA^S83L^* allele during the first ciprofloxacin treatment had only modest effect on the observed resistance (**Figure 3F**). More broadly, species that transiently increased in abundance during treatment likely expanded into niches vacated by more sensitive taxa, with their success governed by competitive release and resource availability. Importantly, prior measurements of viable CFU counts in this system indicate that total biomass often recovers during or shortly after antibiotic treatment^14^, suggesting that these blooms reflect true expansion in absolute abundance rather than purely compositional effects. As disrupted species recovered or went extinct, these ecological conditions shifted, reshaping selective pressures encountered during subsequent treatments. Consequently, identical antibiotic regimens applied to communities with subtly different histories produced divergent outcomes, even among co-housed mice. The extent of this divergence is notable because the large bacterial population size in mammalian guts is often assumed to make adaptive responses deterministic. Instead, our results suggest that ecological context and strain transmission can maintain substantial stochasticity even in large populations. Hence, antibiotic selection in the gut operates on a moving fitness landscape defined by community composition and prior perturbations, linking early ecological events to later evolutionary trajectories.

Ecological openings created by antibiotic perturbation also appeared to enable extensive strain transfer between hosts, revealing transmission as a key determinant of post-antibiotic recovery in our dataset. Using cross-housing experiments between mice colonized with microbiota from two human donors, we observed bidirectional transfer of strains whose timing and success depended on the recovery dynamics of the donor population (**Figure 6**). In most cases, strain transfer occurred after the corresponding resident strain in the recipient mouse declined below the limit of detection, suggesting that antibiotic-induced niche clearance facilitates engraftment of incoming strains. However, we also identified rare instances of strain replacement without prior apparent extinction of the resident strain (**Figure 6G**), indicating that coexistence barriers can sometimes be overcome abruptly and independently of antibiotic-driven collapse^14^. Notably, when transfer occurred, the incoming strain consistently displaced the resident strain rather than coexisting with it, highlighting strong competitive exclusion at the strain level in our system. These dynamics were invisible to species-level profiling and only became apparent through strain-resolved metagenomic analysis. Together, these observations demonstrate that antibiotic exposure not only reshapes within-host evolutionary trajectories but also modulates between-host microbial connectivity, linking recovery outcomes to the timing, directionality, and ecological context of strain transmission. Although opportunities for strain exchange are likely greater in mice due to coprophagy, transmission was far from universal: many disrupted communities did not acquire detectable incoming strains. This observation suggests that constraints on strain transfer and engraftment will be even more pronounced in humans, where opportunities for exchange are more limited.

In addition to selecting on bacterial strains, antibiotic exposure also perturbed the gut microbiome through activation of mobile genetic elements, particularly prophages. We observed stochastic induction of prophages in multiple species (**Figure 7**). These dynamics suggest that phage-mediated lysis can act synergistically with antibiotics to reshape bacterial populations, altering competitive relationships, and accelerating population collapse or replacement. Prophage induction therefore represents an additional axis along which antibiotic perturbations can restructure the microbiome, potentially altering bacterial abundance on faster timescales. Together with strain transfer and genetic sweeps, these results highlight antibiotics as multi-layer perturbations that simultaneously act on bacterial genomes, community structure, and the mobilome, amplifying the potential for divergence in recovery trajectories across hosts.

More broadly, our findings highlight that microbiome recovery from antibiotics is a path-dependent process shaped by transient ecological opportunities, rapid genetic change, host-to-host exchange of strains and phage-associated activities. Even brief antibiotic exposures were sufficient to drive durable genetic sweeps, alter prophage activation, and promote strain replacement rather than restoration of pre-existing populations. Importantly, apparent species-level resilience often masked extensive strain-level turnover, challenging the notion that species-level recovery implies a return to the original community state. These results suggest that effective post-antibiotic interventions, whether aimed at restoring function, limiting resistance, or promoting stable community assembly, must account for the timing and context in which ecological niches open and close. In particular, early introduction of a live biotherapeutic product following an antibiotic exposure may be critical if a closely related resident strain recovers during the antibiotic treatment. In addition to the processes analyzed here, this controlled, densely sampled metagenomic framework provides an opportunity to interrogate other antibiotic-responsive dimensions of microbiome dynamics, including remodeling of metabolic potential, horizontal gene transfer, and broader mobilome implications, offering a path toward a more integrated understanding of how perturbed gut ecosystems are rebuilt, and enable the rational design of interventions that steer post-perturbation trajectories toward more resilient gut communities.

### Limitations of the study

Several features of our experimental and analytical design were critical for resolving these multilayered antibiotic responses, while also defining important limitations. Dense longitudinal sampling enabled us to distinguish transient from durable genetic changes based on shared temporal patterns and to identify selective sweeps that would be missed in sparsely sampled studies. To increase sensitivity, in many strain-level analyses, we aggregated metagenomic reads across mice within cages at each time point, improving detection of low-frequency variants while preserving temporal resolution. This approach highlights a general tradeoff between sequencing depth and cohort size in longitudinal metagenomic studies and suggests that structured aggregation can be an effective strategy when biological replication is available. At the same time, our analyses were constrained by the resolution limits of short-read sequencing. Structural variants, local amplifications, mobile genetic elements, and recombination events that may underlie resistance or susceptibility phenotypes can be difficult to resolve without long-read or assembly-based approaches. These limitations extend to prophage analyses: while we detected antibiotic-associated phage dynamics and inferred induction events from temporal abundance patterns, distinguishing integrated from excised prophages or resolving circularized genomes and free virions from short-read data is challenging. More definitive characterization of phage induction, host linkage, and horizontal gene transfer will require complementary approaches such as long-read sequencing, viral particle enrichment, or proximity ligation strategies^19,44,45^. In addition, phage and bacterial abundances were inferred through reference-based classification, and incomplete representation of viral diversity may limit sensitivity for detecting low-abundance or novel phages. Thus, some antibiotic-responsive mobilome dynamics may remain unresolved. Finally, while humanized mouse models provide experimental control and reproducibility that are not feasible in human studies, differences in housing, exposure, and transmission dynamics (such as the increased opportunities for microbial exchange due to mouse coprophagy) must be considered when extrapolating these findings to human populations. Together, these considerations emphasize both the strengths of this framework for dissecting causal mechanisms and the need for complementary approaches to fully capture the complexity of antibiotic-microbiome interactions.

## RESOURCE AVAILABILITY

### Lead contact

Requests for further information and resources should be directed to and will be fulfilled by the lead contact, Kerwyn Casey Huang (^10^).

### Materials availability

This study did not generate unique reagents.

### Data and code availability

- Shotgun metagenomic sequencing data have been deposited at NCBI SRA as BioProject number PRJNA1467299.
- All original code has been deposited to https://bitbucket.org/kchuanglab/mouse-abx-metagenomics/.
- Any additional information required to reanalyze the data reported in this paper is available from the lead contact upon request.

## ACKNOWLEDGEMENTS

We thank the members of the Huang and Good labs for helpful discussions. We would like to thank Stanford University and the Stanford Research Computing Center for providing computational resources and support that contributed to this research, some of which was performed on the Sherlock cluster. This work was funded by a Stanford PRISM Baker Fellowship (to J.A.L.), a James S. McDonnell Postdoctoral Fellowship (to H.S.), Human Frontier Science Program grant RGEC33/2023 (to B.H.G.), NSF award EF-2125383 (to K.C.H.), and NIH awards R01 DK085025 (to J.L.S.), R35 GM146949 (to B.H.G.), R01 AI147023 (to K.C.H.), RM1 GM135102 (to K.C.H.), and DP1 DK147449 (to. K.C.H.),. K.C.H., B.H.G., and J.L.S. are Chan Zuckerberg Biohub Investigators.

## AUTHOR CONTRIBUTIONS

Conceptualization, C.L., M.K., E.Y., J.A.L., F.B.Y., K.N., J.L.S., B.H.G., K.C.H., and H.S.; methodology, C.L., M.K., F.B.Y., K.N., and H.S.; software, C.L., M.K., J.A.L., and H.S.; validation, C.L., M.K., and H.S.; formal analysis, C.L., M.K., and H.S.; investigation, C.L., M.K., E.Y., J.A.L., F.B.Y., K.N., J.L.S., B.H.G., K.C.H., and H.S.; resources, C.L., M.K., F.B.Y., K.N., J.L.S., B.H.G., K.C.H., and H.S.; data curation, C.L., M.K., K.N., and H.S.; writing – original draft, C.L., M.K., B.H.G., K.C.H., and H.S.; writing – review & editing, C.L., M.K., E.Y., J.A.L., F.B.Y., K.N., J.L.S., B.H.G., K.C.H., and H.S.; visualization, C.L., M.K., and H.S.; supervision, J.L.S., B.H.G., K.C.H., and H.S.; project administration, B.H.G., K.C.H., and H.S.; funding acquisition, J.L.S., B.H.G., and K.C.H..

## SUPPLEMENTARY FIGURES AND FIGURE LEGENDS

**Figure S1:**
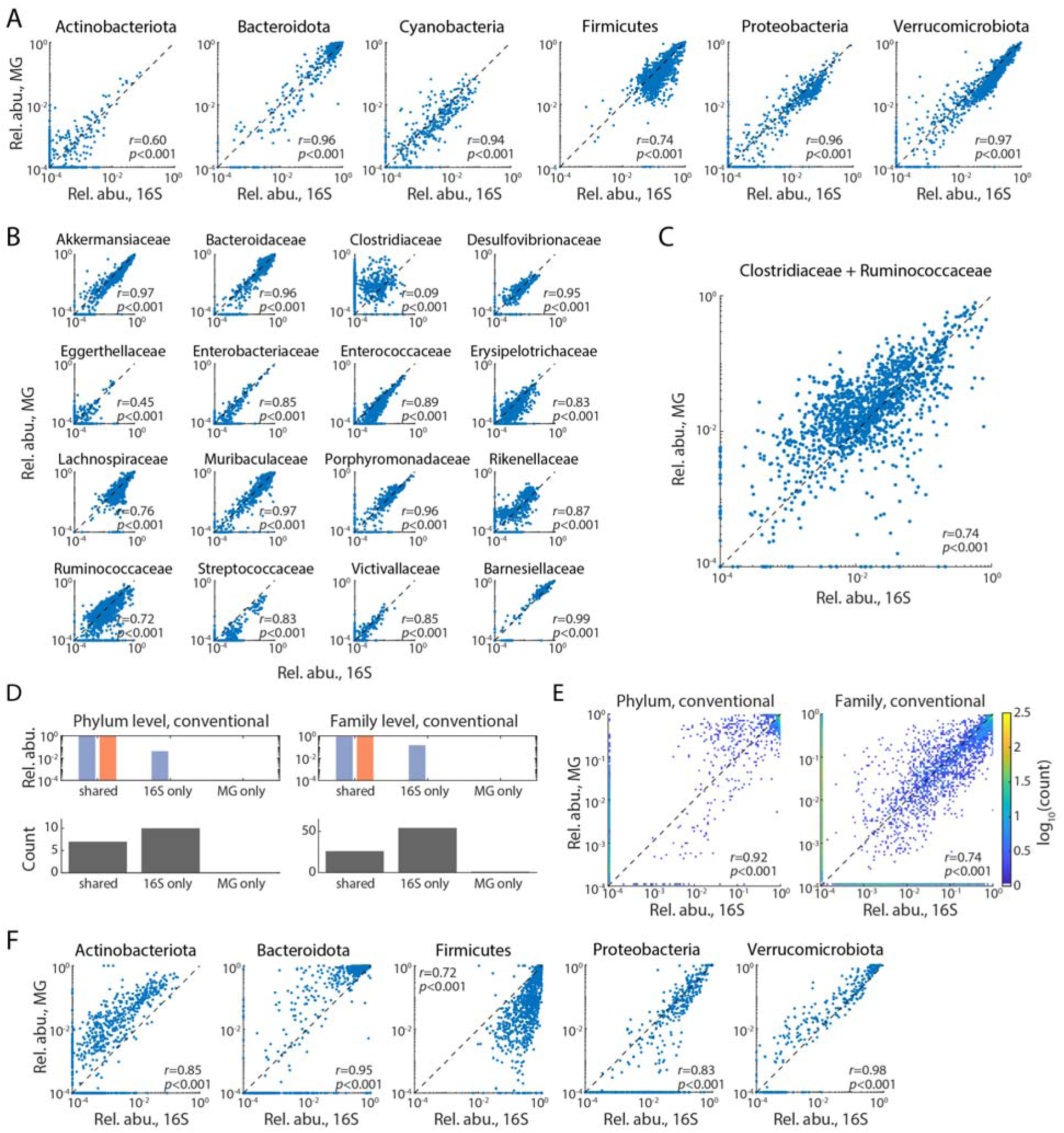
Metagenomic sequencing recapitulates phylum- and family-level ecological dynamics observed by 16S result profiling, related to. Figure 1. (A,B) Scatter plots comparing relative abundances estimated by metagenomic (MG) sequencing and 16S rRNA profiling for taxa at the phylum (A) and family (B) levels in humanized mouse fecal samples (prevalence ≥10%). No systematic deviations are observed at the phylum level. At the family level, correlations are generally strong, although some low-abundance families show increased variability, likely reflecting detection noise. Metagenomic estimates tend to overestimate *Clostridiaceae* and underestimate *Ruminococcaceae* abundances. (C) Scatter plot of the combined relative abundance of *Clostridiaceae* and *Ruminococcaceae*. The summed abundance shows strong agreement between metagenomics and 16S, suggesting that discrepancies between these families arise from differences in taxonomic assignments across reference databases. (D) Metagenomics fails to detect a larger fraction of taxa in conventional mice compared to humanized mice. Across 1374 fecal samples from 104 mice, analysis of metagenomics data identified 7 of 17 phyla and 26 of 80 families detected by 16S profiling, accounting for ∼95.9% and 85.7% of total abundance. (E) Conventional mice exhibited lower concordance between metagenomics- and 16S- based relative abundance estimates than humanized mice. (F) Scatter plots comparing phylum-level relative abundances (prevalence ≥10%) in conventional mice. Metagenomic estimates systematically underestimate the abundance of Firmicutes, but overall abundances remain well correlated with 16S measurements.

**Figure S2:**
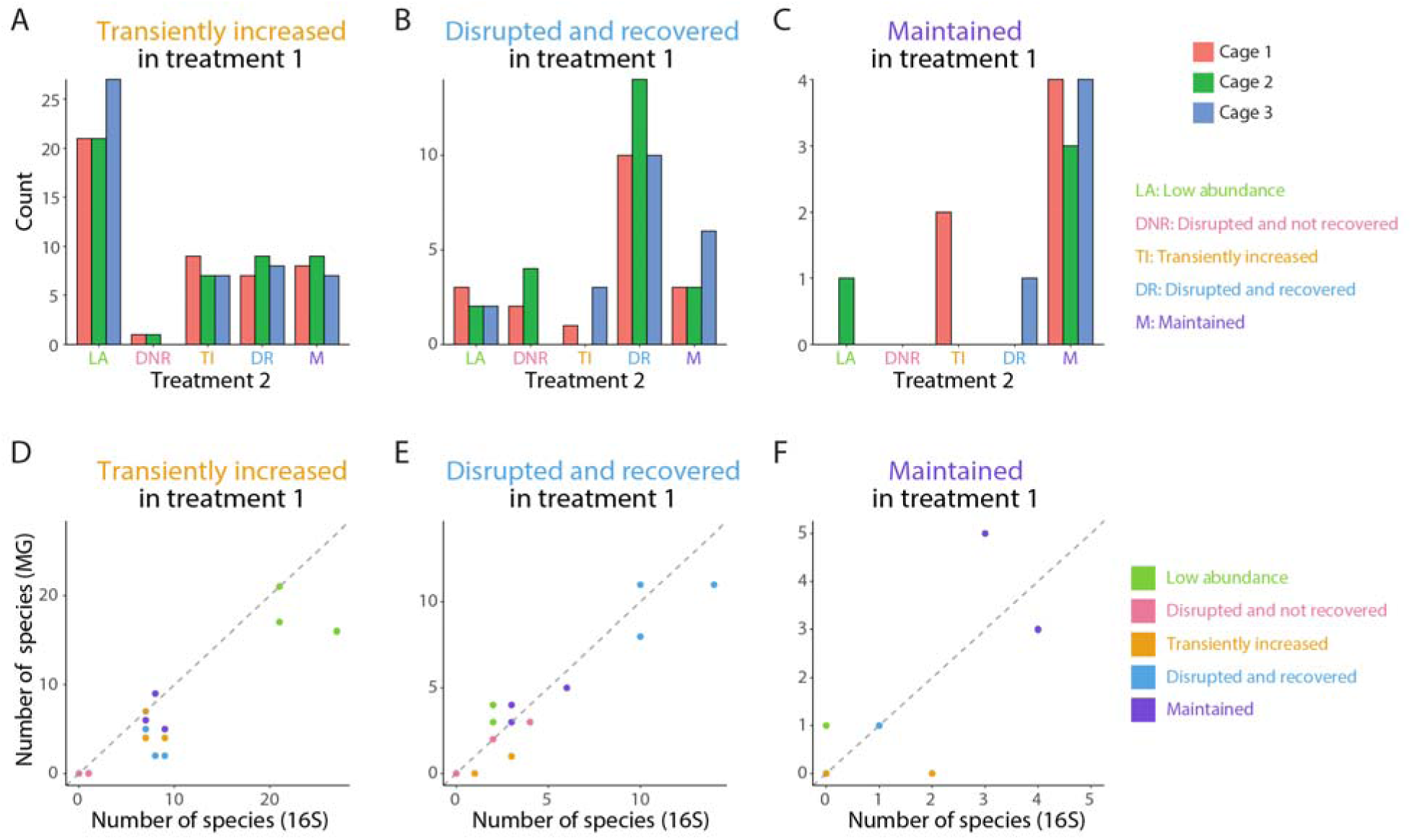
Metagenomic and 16S profiling yield concordant classifications of species responses to repeated ciprofloxacin treatment, related to Figure 2. (A–C) Bar plots showing the distribution of species response classes during the second treatment, stratified by their response during the first treatment, based on 16S rRNA profiling. Species responses were broadly consistent across cages. (D–F) Comparison of species response classifications during the second treatment between 16S and metagenomic profiling, stratified by first-treatment responses. Each point represents the number of species assigned to a given response class by the two methods. Classifications are highly concordant between approaches, indicating consistent quantification of species-level dynamics. In several cases, metagenomic profiling identified additional species within a response class, reflecting increased sensitivity.

**Figure S3:**
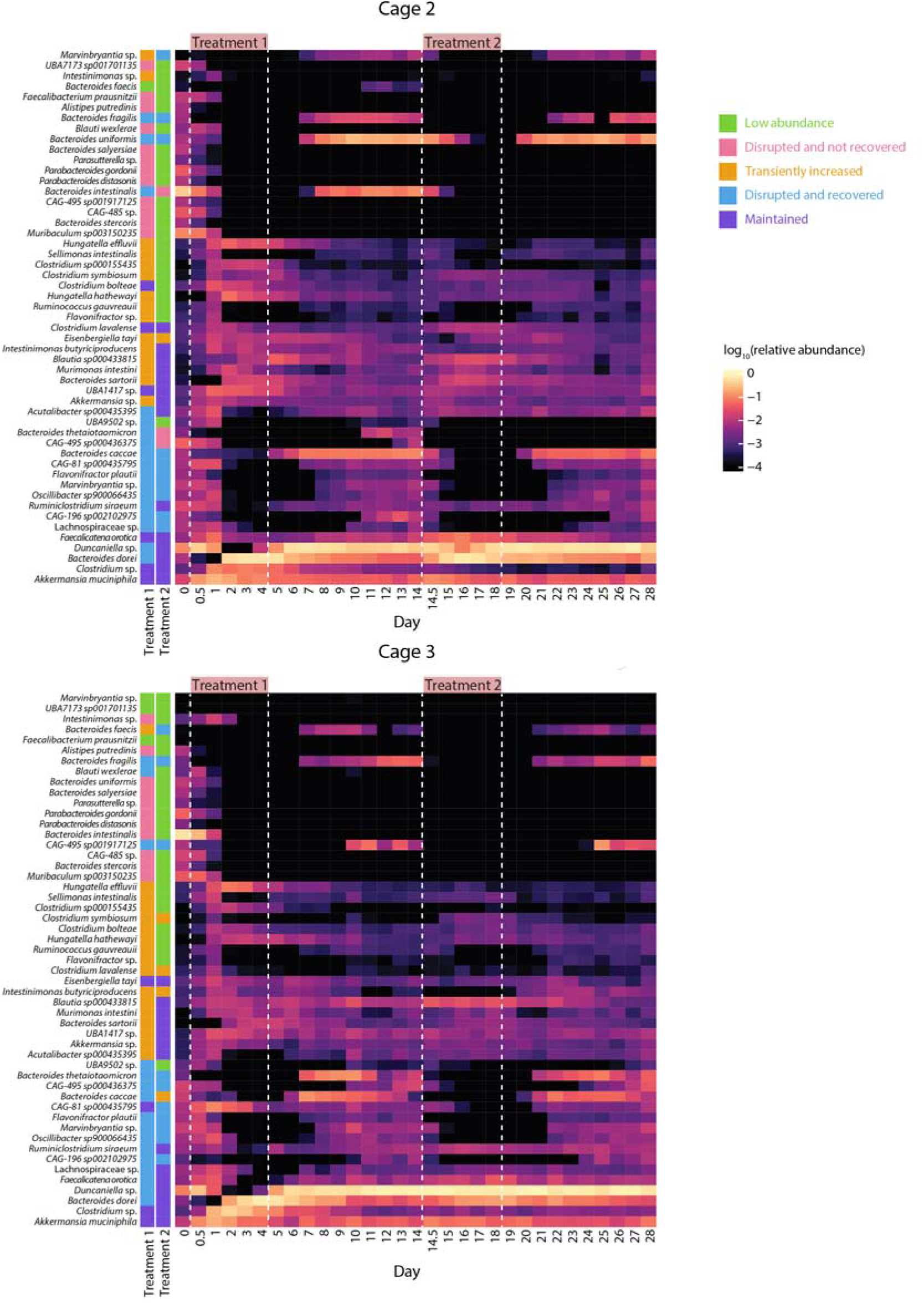
Species-level responses to repeated ciprofloxacin exposure are broadly reproducible across cages, related to Figure 2. (A,B) Heatmaps showing relative abundances of high-abundance species (maximum abundance >1%) in cages 2 (A) and 3 (B) during the double ciprofloxacin treatment in humanized mice. Rows represent species and columns represent time points, with treatment periods indicated above. Species are grouped by response class (left annotation). Species-level responses to ciprofloxacin are largely consistent across cages, with similar patterns of disruption, recovery, and persistence observed despite independent housing. Vertical dashed lines denote antibiotic treatment windows.

**Figure S4:**
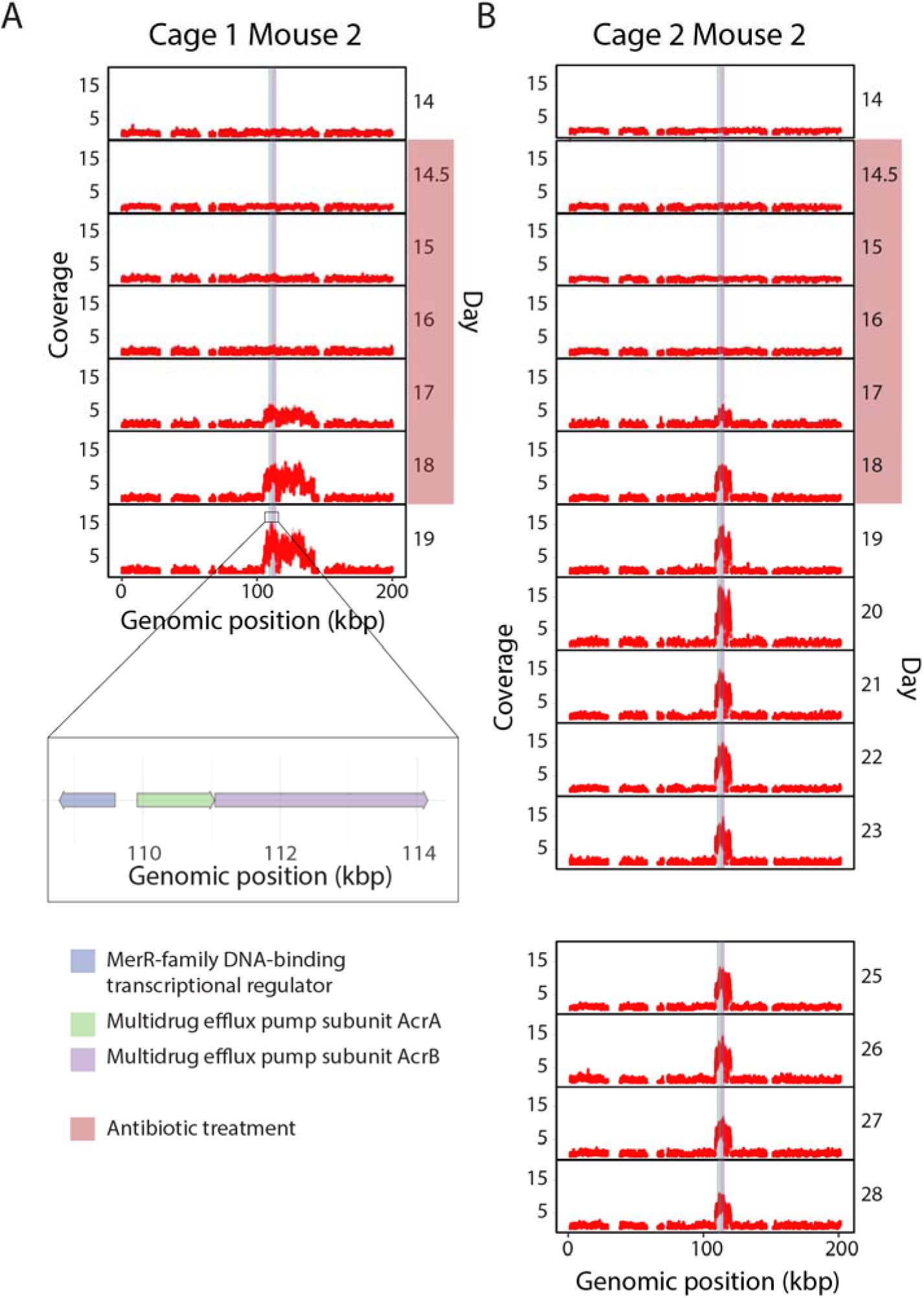
Amplification of multidrug resistance-associated loci provides an alternative route to ciprofloxacin resistance in *A. muciniphila*, related to Figure 4. (A-B) Relative coverage profiles across the *A. muciniphila* genome (0–200 kbp) in two mice during the second ciprofloxacin treatment. Points denote individual genomic positions, highlighting putative amplification events. These two mice along with the mouse shown in Figure 4 are the only individuals exhibiting amplification of this region, which contains three genes tightly linked to ciprofloxacin resistance. Amplification emerged around day 17, during the second treatment, and persisted through the remainder of the experiment. Co-amplification of the MerR regulator and adjacent efflux pump genes suggests a coordinated genetic mechanism that may contribute to reduced antibiotic susceptibility. Time points lacking data correspond to low-abundance samples with insufficient coverage for reliable SNV detection. Mouse 2 in cage 1 was sacrificed at day 19, and therefore no data are available thereafter.

**Figure S5:**
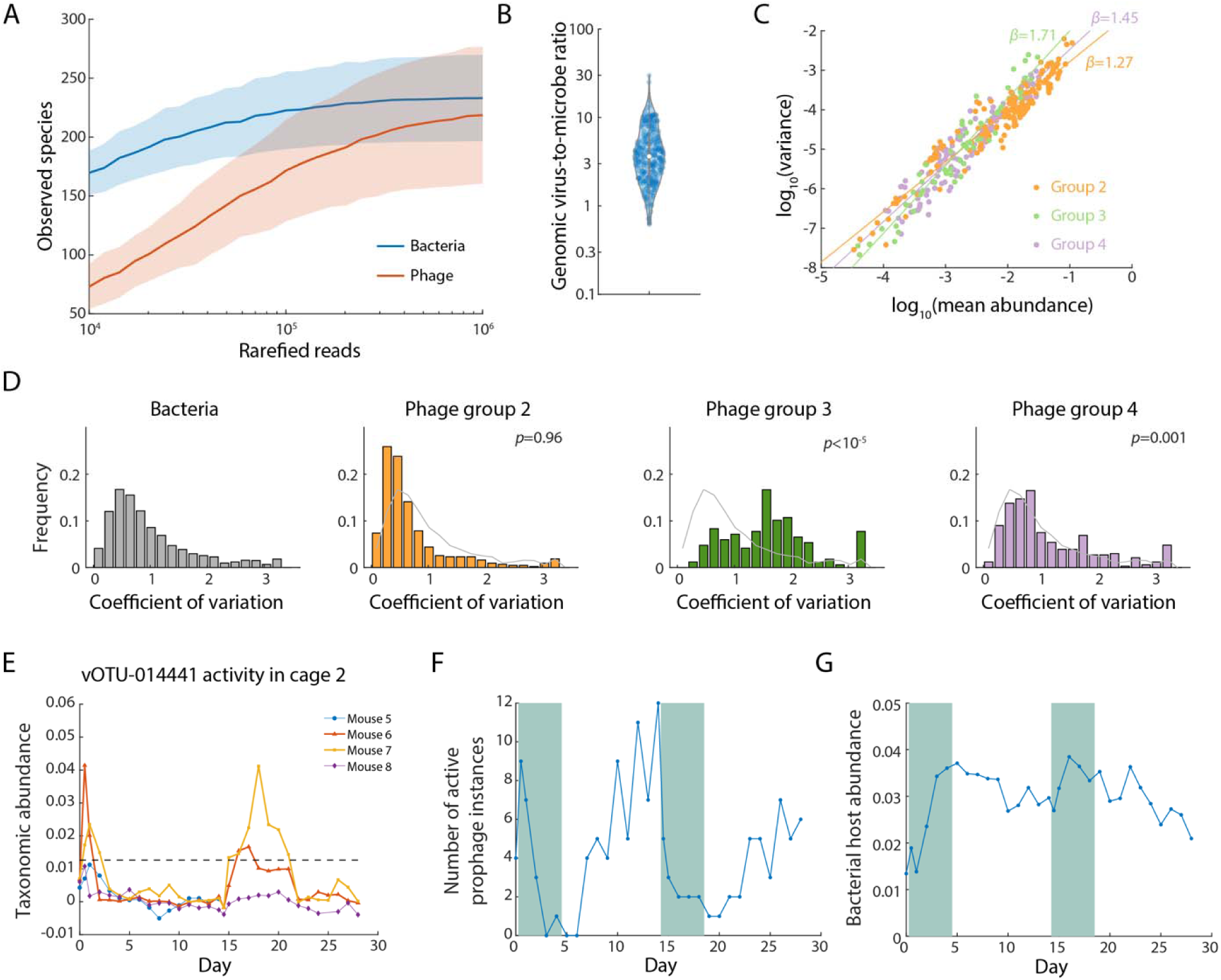
Phage populations in humanized mice exhibit heterogeneous temporal dynamics, related to Figure 7. (A) Rarefaction analysis of bacterial and phage diversity in day 0 samples from the two-course ciprofloxacin experiment in humanized mice. Phage profiles show a steeper rarefaction curve than bacteria, consistent with their smaller genome sizes. (B) Distribution of virus-to-microbe ratios across all samples, calculated from Phanta-estimated viral and bacterial abundances. (C) Taylor’s law analysis relating temporal variance to mean abundance for phages in groups II, III, and IV. All three groups exhibit power-law scaling, with group III showing the largest exponent and group IV an intermediate exponent, consistent with their increasingly variable dynamics. (D) Inter-mouse heterogeneity of bacteria and phages, quantified as the coefficient of variation (CV) across mice at each time point. Group II phages show CV distributions comparable to those of bacterial hosts, whereas groups III and IV exhibit elevated variability across mice. (E) Phage activity for another vOTU in cage 2, in which activity was detected in only two mice (mice 6 and 7). Dashed line: threshold for calling phage activity for this vOTU. Mouse 5 was sacrificed on Day 14. (F) Total number of group IV phage activity events detected at each time point across the experiment. Phage activity did not increase during ciprofloxacin treatment. (G) Abundances of bacterial hosts corresponding to group IV phages. Host bacterial abundances do not show parallel changes that would account for the reduction in phage activity during ciprofloxacin treatment.

## SUPPLEMENTARY TABLES

**Table S1: Summary of all mouse experiments in ref.** ^14^. All mouse fecal samples were sequenced and analyzed in this study via metagenomics. Day 0 denotes the start of the first antibiotic treatment.

**Table S2: Cross-referenced curation of taxonomic annotations between the GreenGenes and UHGG databases used for comparative analyses.**

## METHODS

## EXPERIMENTAL MODEL

### Sample collection

Mouse fecal samples were collected as described previously^14^.

## METHOD DETAILS

### Library preparation and sequencing

Genomic DNA from whole fecal pellets were extracted using PowerSoil and PowerSoil-htp kits (Qiagen) as described previously^14^, with their concentrations quantified in 384-well format using the Quant-iT PicoGreen dsDNA Assay Kit (ThermoFisher).

Sequencing libraries were generated in 384-well format with a custom low-volume protocol adapted from the Nextera XT process (Illumina) as described previously^46^. Briefly, each genomic DNA sample was normalized to 0.18 ng/µL, and tagmentation, neutralization, and PCR steps of the Nextera XT protocol were performed on a Mosquito HTS liquid handler (TTP Labtech). This low-volume protocol used 0.072 ng of genomic DNA input, 11 PCR cycles, and yielded a final volume of 4 µL per library.

Custom 12-bp dual unique indices were used as barcodes. The resulting libraries were pooled and cleaned up using AMPure XP beads (Beckman), targeting a fragment length of 450 bp (insert size ∼350 bp).

Sequencing was performed using NovaSeq 6000 on S4 flow cells in the 2×150 bp configuration at Chan Zuckerberg Biohub, targeting 10 million paired-end reads per sample. A total of 45 G paired reads, ∼13 terra base pairs of metagenomic data were generated.

### Metagenome quality control and pre-processing

Raw sequencing reads were demultiplexed and processed using BBtools suite^47^: exact duplicate reads were removed with the clumpify command, and adapters and low-quality bases were trimmed (bbduk command; trimq=16, minlen=55). The trimmed reads were mapped using the BBmap command against the mouse genome to remove host reads. The paired reads were then merged (BBMerge; rem k=62 extend2=50 ecct vstrict)^48^. After the process, a median of 9 M high-quality, non-duplicated reads were retained for each sample.

### MIDAS2 species profiling

To increase coverage of metagenomic sequencing for certain analyses, FASTQ files from all fecal samples in the same cage at a given time point were concatenated together. The MIDAS2 pipeline described below was applied to both individual mouse fecal samples and these cage-level concatenated datasets.

Paired-end metagenomic FASTQ files were processed with the MIDAS2 pipeline with default parameters^29^, profiling species and quantifying relative abundances from coverage across 15 universal, single-copy marker genes. Reads with alignment coverage below 0.75 were excluded, and only marker genes supported by at least two reads were retained. Species were only reported if at least two marker genes were detected. Relative abundance estimates were derived from the normalized coverage of each species’ marker genes. Species- and gene-level annotations were referenced against the UHGG (Unified Human Gastrointestinal Genome) database^49^.

### MIDAS2 SNV profiling

SNV calling on metagenomic data from individual mice and combined cages at each time point was performed using MIDAS2. SNVs were called for all species in each sample with median marker coverage greater than 2, discarding reads with alignment identity less than 94, mean quality less than 20, or alignment coverage less than 0.75, as well as discarding bases with quality less than 30. Population SNVs for each sample were computed for each bacterial species, retaining sites in the genome with more than two post-filtered reads. Default MIDAS2 parameters in source code related to SNV calling, including minimum allele frequency thresholds, were relaxed to capture all candidate variants, with downstream filtering applied during post-processing.

### Profiling phage and bacteria abundances using Phanta

Phanta was run with default parameters, including a confidence threshold of 0.1 (“confidence_threshold”), viral genome coverage threshold of 0.1 (“cov_thresh_viral”), viral unique minimizer threshold of 0 (“minimizer_thresh_viral”), bacterial genome coverage threshold of 0.01 (“cov_thresh_bacterial”), and bacterial unique minimizer threshold of 0 (“minimizer_thresh_bacterial”).

## QUANTIFICATION AND STATISTICAL ANALYSIS

### Comparison between 16S and metagenomic datasets

Previous 16S sequencing data were retrieved from Ref. ^14^, using annotations from GreenGenes^50^. The MIDAS2 profiling used the Unified Human Gastrointestinal Genome (UHGG) database^30^. The different taxonomic annotations between the two databases were manually curated (**Table S2**) at the family level and higher. Since GreenGenes lacks many annotations at genus or species level, we did not attempt to directly compare the two profiling methods at these levels.

### Species response classification

For each antibiotic treatment, cage- and mouse-level species responses were classified into five types of behaviors using their relative abundance dynamics. A species was classified as maintained if it had an initial abundance greater than 10^-3^ immediately before antibiotic treatment, as well as either a minimum abundance during treatment greater than 10^-3^ or minimum abundance during the treatment exceeding one-tenth of the starting abundance. A species was classified as disrupted if the abundance at the time point immediately before antibiotic treatment was greater than 10^-3^, and the minimum abundance during treatment was both less than one-tenth of this starting abundance and less than 10^-3^. Disrupted species that had post-antibiotic abundance exceeding one-tenth of the starting abundance were classified as *Disrupted and recovered*, and species that remained less than one-tenth of the starting abundance post-antibiotics were classified as *Disrupted and not recovered*. Species with starting abundance less than 10^-3^ immediately before antibiotic treatment were classified as *Low abundance*. Low-abundance species that had maximum abundance during or after treatment greater than 10^-3^and exceeding ten times the starting abundance were classified as *Transiently increased*. All species were able to be classified into these five categories.

### SNV sweep detection

Prior to detecting SNV sweeps in each species, several filtering steps were performed. Reference and alternate alleles for each position were arbitrarily selected as the most and second most frequent allele at that position across all samples, respectively. Tri- and quad-allelic sites were treated as bi-allelic using the two most frequent alleles across the population. Only protein-coding sites were considered for this analysis. Mutations were classified as synonymous or nonsynonymous.

MIDAS2 outputs the sequencing coverage *D_it_* for a site *i* at a time point *t* for each high-abundance bacterial species in a single sample. The median coverage across all protein-coding sites with more than two reads in a species,*D_t_*, was calculated from this. For a given species, time points with *D_t_* below 10 were excluded from further analysis. Sites with abnormally small or large coverage were filtered to reduce mismappings from the database, masking sites at each time point with *D_it_* < *D_t_* /3 or *D_it_* > 3*D_t_* . Sites that were masked in more than 25% of time points were excluded.

Allele frequency at a site was estimated by dividing the measured allele counts by the total coverage at that site. SNVs that had both minor allele frequency (MAF) less than 20% in at least one time point and greater than 80% at another time point were classified as sweeping.

### Clustering of SNV trajectories

Sweeping SNVs were clustered based on their temporal behavior across individual mice and cages. The goal of this approach was to visualize consistent SNV patterns and replicability across the experiment. To do this, a custom -means clustering approach was developed.

We defined a distance function that incorporates expected binomial sampling variance in allele frequency estimates. For a given SNV *_i_*, let *f_itc_* represent the alternate allele frequency at time point *t* in cage (or mouse)*c*, and let *D_itc_* be the corresponding total read coverage. For a pair of trajectories *f_i_* and *f_j_*, we computed the pairwise distance:

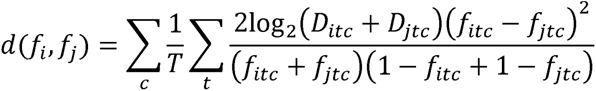

However, the relative orientation of SNVs in the population is unknown, and thus the pairwise distance is calculated with respect to both the reference and alternate allele, referred to as polarizations:

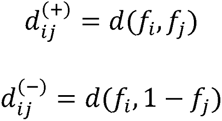

The effective distance between a pair of trajectories was defined as the smaller of the positive and negative polarizations:

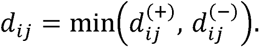

When 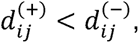, the two SNVs were considered to have consistent polarization; otherwise, their trajectories were treated as flipped in orientation. This distance formulation penalizes discrepancies in allele frequency while down-weighting differences that can be explained by limited sequencing depth through the logarithmic coverage term. Pairs of trajectories with identical temporal patterns yield *di_j_*_, ≈ 0,_ whereas unrelated or oppositely polarized trajectories produce larger values.

*k*-means clustering was performed on sweeping SNV trajectories across mice and cages, with centroid initialization following a modified *k*-means++^51^ approach to ensure ^51^that the starting *k* centroids were well separated across the allele-frequency space. The first centroid (allele frequency trajectory) for *k*-means clustering was chosen at random from the list of sweeping SNVs. For each remaining site, distance to the nearest existing centroid was computed using the previously described distance metric. New centroids were then iteratively sampled with probability proportional to the squared distance from the nearest centroid, producing a diverse set of cluster initializations.

Clustering was then performed, with SNVs assigned to the nearest centroid based on the minimum distance across both polarities. Centroids were updated in each iteration with appropriately polarized trajectories. This process was repeated until convergence or until the maximum number of iterations (10) was reached.

The number of clusters (*k*) was evaluated from 2 to 10, and the optimal *k* was chosen based on a combination of the average Silhouette score (computed from pairwise distances under the custom metric) and visual inspection of cluster coherence.

After clustering, centroids and their associated SNVs were reoriented to ensure consistent polarity across clusters. A global reference centroid was selected as the centroid with the smallest total distance to all centroids. Each remaining centroid was then flipped, if necessary, to minimize its distance to this reference. Finally, all SNV trajectories within a cluster were subsequently adjusted to match the final polarity of their respective centroid.

SNVs were classified based on the number of co-occurring passenger sweeps. SNVs with fewer than three passenger sweeps were classified as *de novo* mutations, while SNVs with larger numbers of associated sweeps were interpreted as arising from pre- existing variation. In practice, SNVs were readily distinguishable between these categories, with few cases near the threshold.

### Detecting regions of focal genomic amplification

To identify regions of genomic amplification, the relative coverage ratio at each site was calculated as the site-specific read depth normalized by the median coverage across all genomic sites in the sample *Dit*/*Dt* 1Genomic regions of high relative coverage ratio, *Dit*/*Dt* >3, were quantified and visualized temporally.

### Identification of MerR family DNA-binding transcriptional regulator

We identified the hypothetical protein in *Akkermansia muciniphila* to have high homology with MerR family DNA-binding transcriptional regulator using NCBI BLASTp^52^ with the NCBI non-redundant (nr) protein sequences database (96.4 percent identify, E value 0.0). Conserved domains were identified using NCBI CD-search^53^ and HMMER^54^. A 3D model was generated with ColabFold^55^, which implements AlphaFold2^56^. The predicted structure was visualized in UCSF Chimera^57^.

### Inference of variable sites in *Akkermansia muciniphila* genome

We filtered sites from the MIDAS2 SNV output to retain only those with greater than 10x total coverage. At each retained position, the most common allele in the metagenome of a given mouse was designated as the major allele. This allele was then used to reconstruct the genome of the dominant strain in that mouse. Across mice, sites were labeled variable if the major allele at that position did not agree across all mice. We defined a consensus allele at each position as the major allele in more than five of the ten mice. For each mouse and each variable site, we then calculated the fraction of sequencing reads supporting this consensus allele by dividing the read count for the consensus nucleotide by the total read depth at that position. Hierarchical clustering using Euclidean distance and complete linkage was performed on consensus allele fractions, clustering both variable sites and strains, with strains clustered separately within ciprofloxacin susceptibility groups.

### Strain transfer during cross-housing

The previously mentioned species and SNV profiling MIDAS2 pipeline was also used to process metagenomic sequences from the cross-housing experiment.

Species transfer was defined as the appearance of a bacterial species in co-housed mice that had previously been unique to one donor group. Specifically, a species was considered transferred if it was present in any separately co-housed mice from H1, absent in all separately co-housed mice from H2 and in cross-housed H2 mice prior to cross-housing, and subsequently detected in cross-housed H2 mice after cross-housing. The same criterion was applied reciprocally for species originating from H2. To distinguish whether the transferred strain matched the donor strain, we first defined a set of “conserved SNVs” by identifying sites where a specific allele was present at >80% frequency in all separately co-housed mice from the single-donor group. We then examined these conserved SNVs in the cross-housed mice: if more than 0.1% of these sites fell below 20% allele frequency, the strain was classified as distinct from the donor strain.

To identify potential strain transfer events, donor-specific "marker SNVs" were defined for each high-abundance bacterial species. Marker SNVs were sites where a specific allele was consistently present at greater than 80% frequency in all separately co-housed mice from one donor group and less than 20% frequency in all separately co-housed mice from the other donor. Hundreds of marker SNVs were typically observed per species. The detection of these donor-specific SNVs in mice from the opposite donor group after cross-housing was interpreted as evidence of strain transfer.

### Phage classification and activity analysis

We focused phage analyses on viral taxa with a median taxonomic abundance of at least 10^−3^. For each phage, we first quantified abundance variation across all samples. Phages whose abundances varied by less than 3–fold were classified as group I. For the remaining phages, we identified bacterial species whose abundances were correlated with phage abundance across all mice and time points (Pearson’s *r*>0.6). Phages for which no correlated bacterial species were identified were classified as group III.

When multiple bacterial species exhibited strong correlations with a given phage, we considered those with correlation coefficients within 0.1 of the maximum value for that phage (*r*>*r*_max_–0.1, where *r*_max_ is the highest correlation observed across all bacteria). Among these candidates, we computed the linear regression slope between bacterial and phage abundances and selected the bacterial species with a slope closest to 1 as the putative host. All inferred phage–host pairs were subsequently visually inspected.

For phages with an identifiable host, residual phage abundance was calculated by fitting a linear model relating phage and host abundances, scaling host abundance accordingly, and subtracting the fitted host-associated component from the observed phage abundance. A phage was considered to exhibit activity beyond host-tracking dynamics if it showed three or more instances (where one instance corresponds to a single mouse at a single time point) in which residual abundance exceeded a threshold defined as median+3×MAD (median absolute deviation). Phages meeting this criterion were classified as group IV, whereas the remaining host-associated phages were classified as group II. For group IV phages, individual instances with residual abundance above this threshold were defined as phage activity events.

