## Supplementary figures and images for "Path-dependent recovery of the gut microbiome after antibiotics emerges from coupled ecological and evolutionary dynamics"

### Figure S1

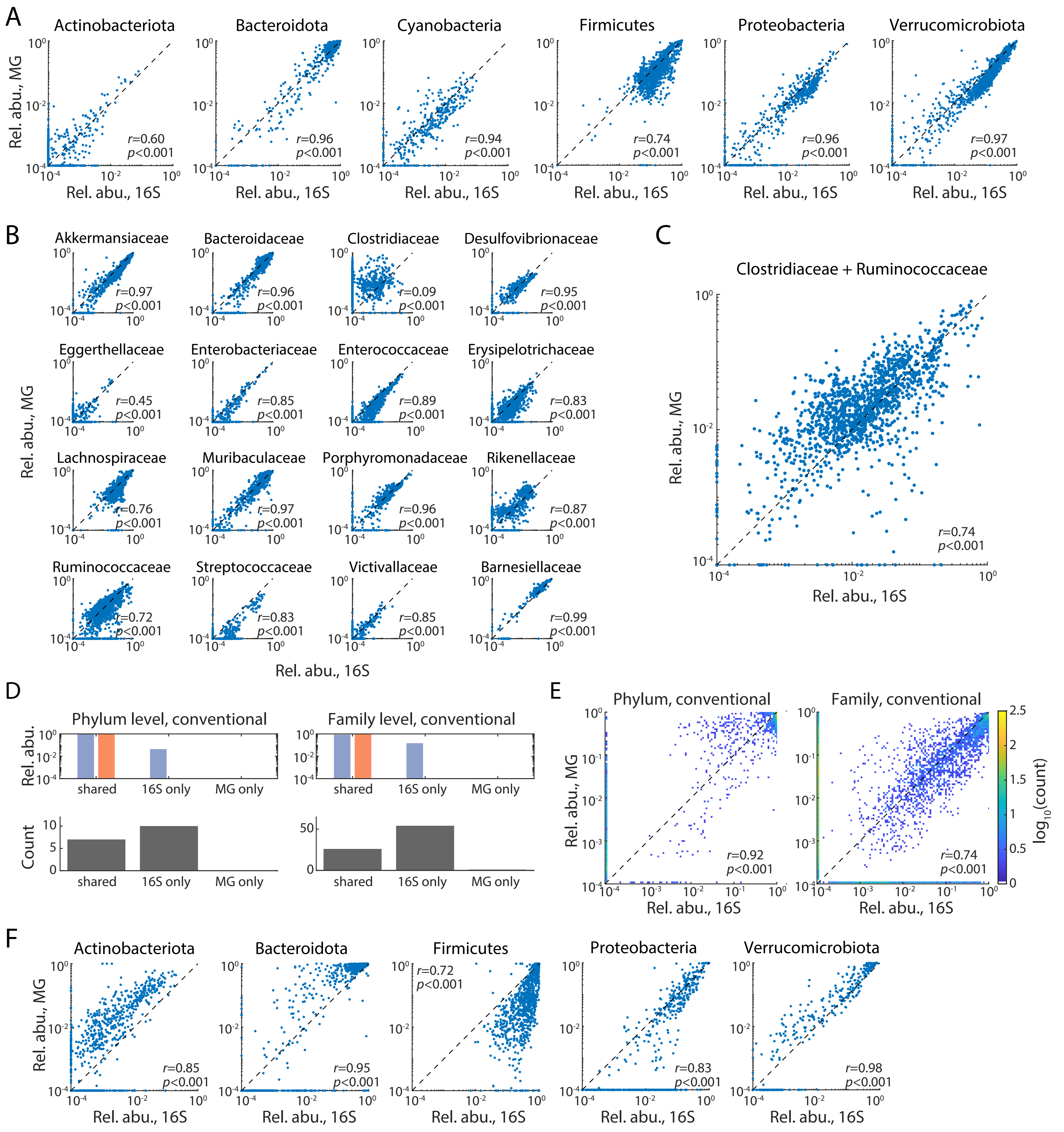

### Figure S2

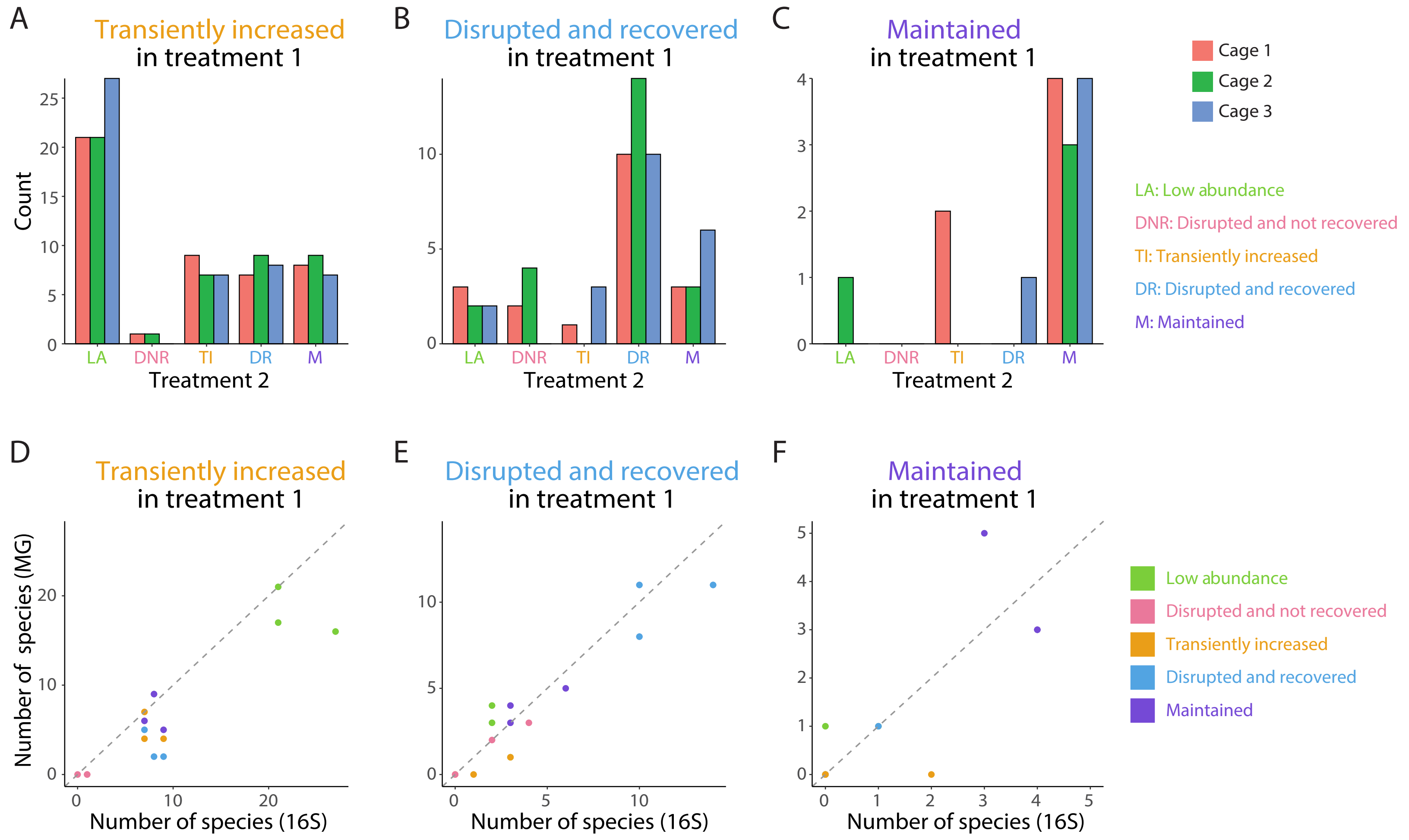

### Figure S3

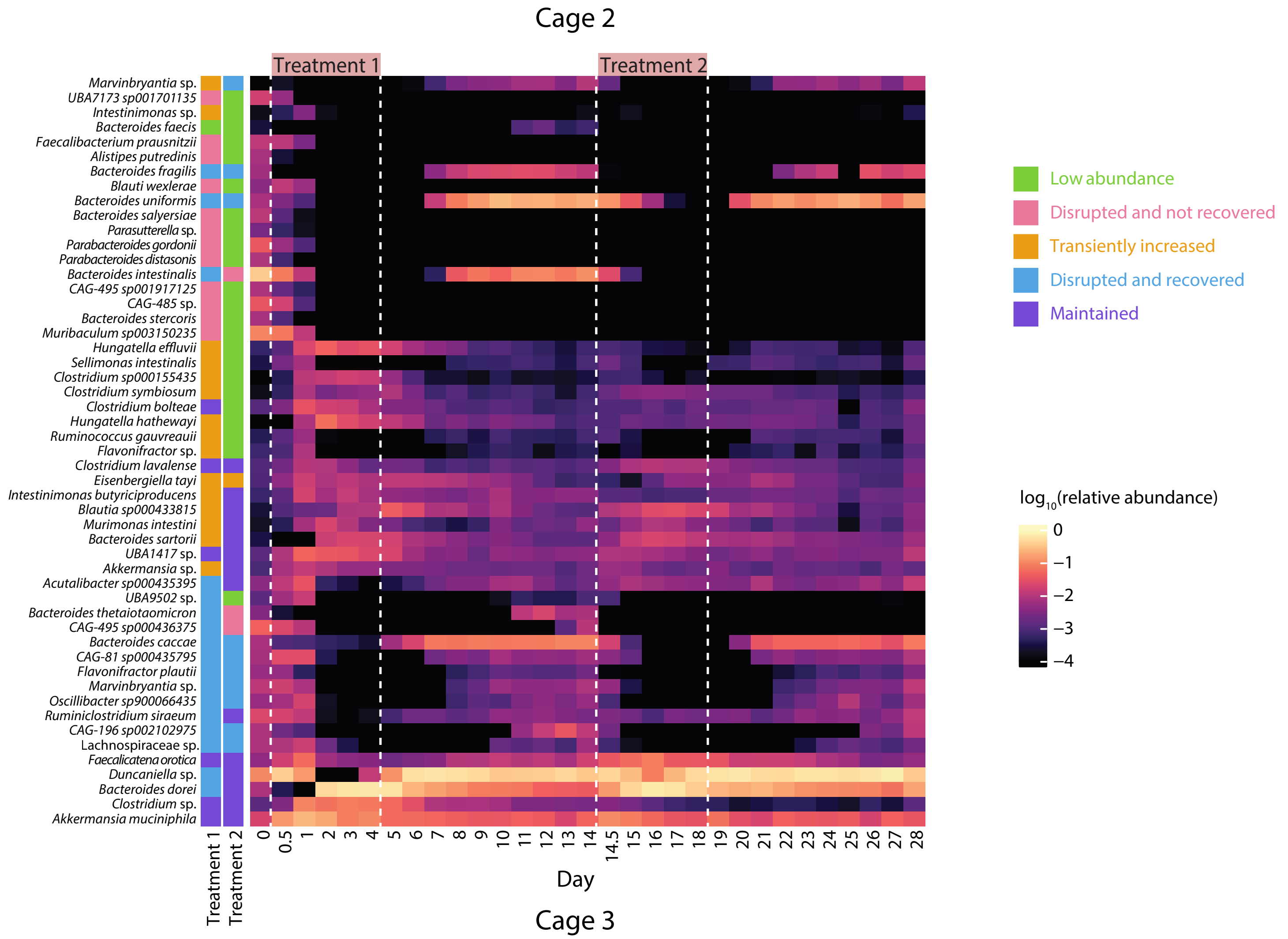

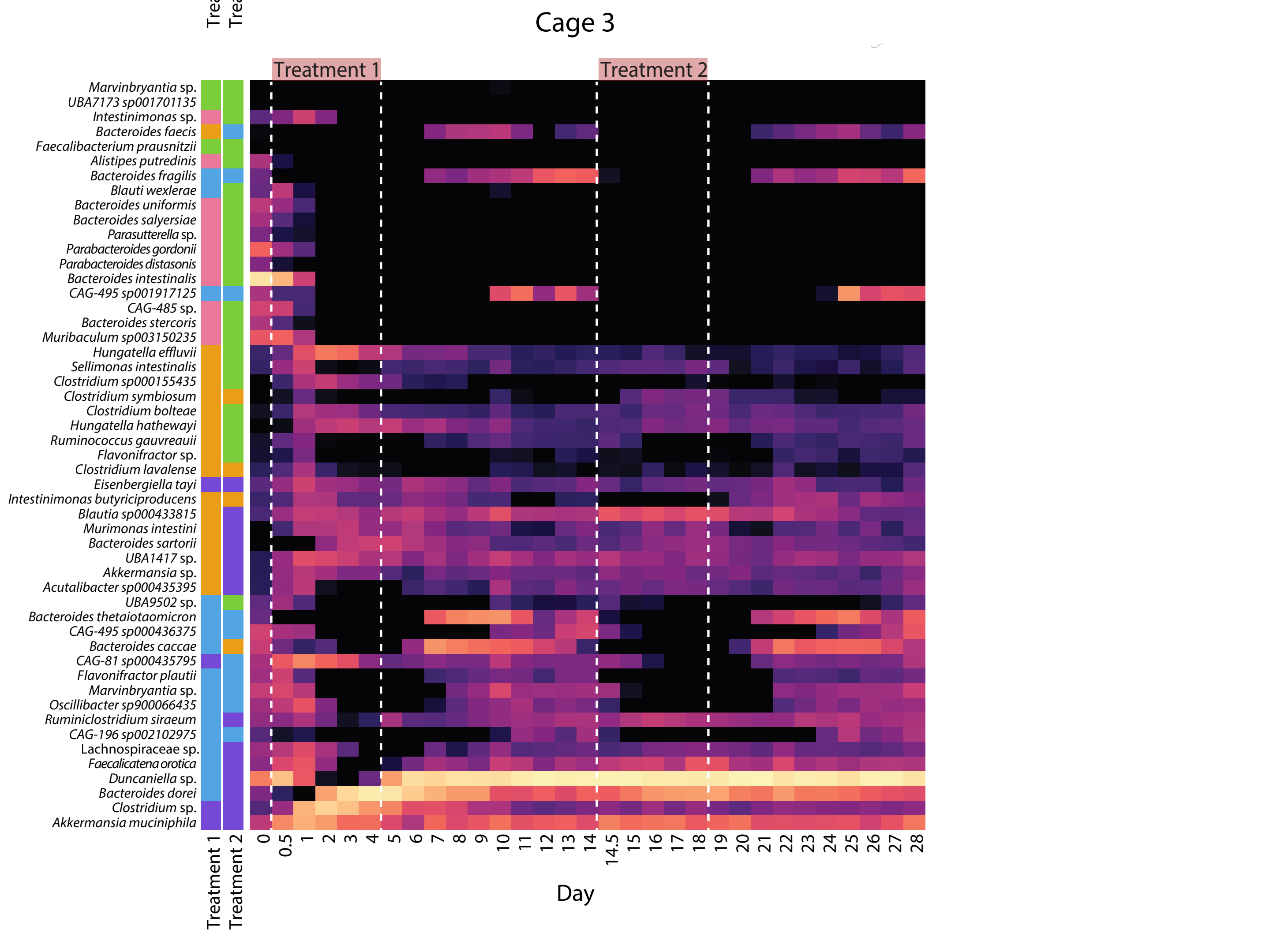

### Figure S4

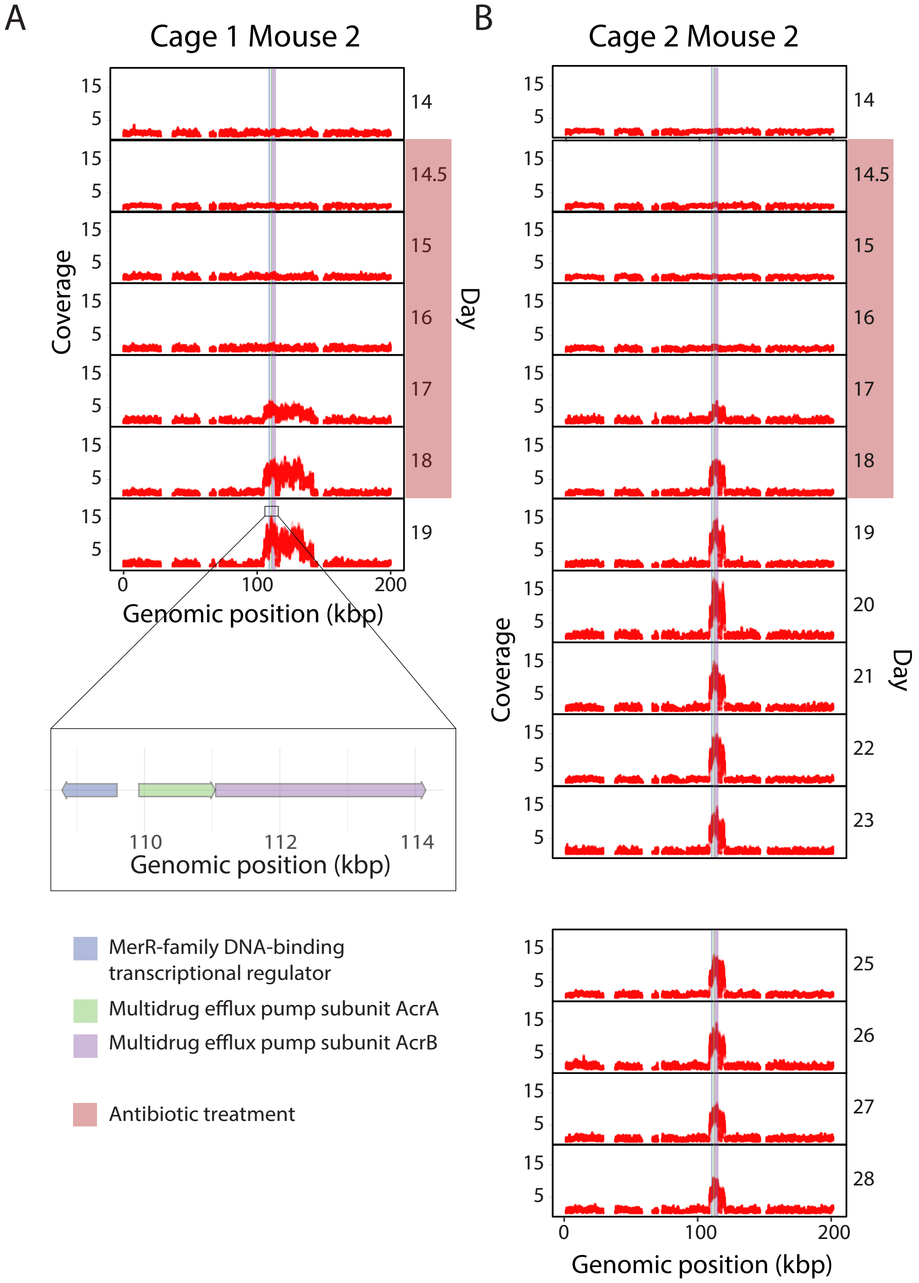

### Figure S5

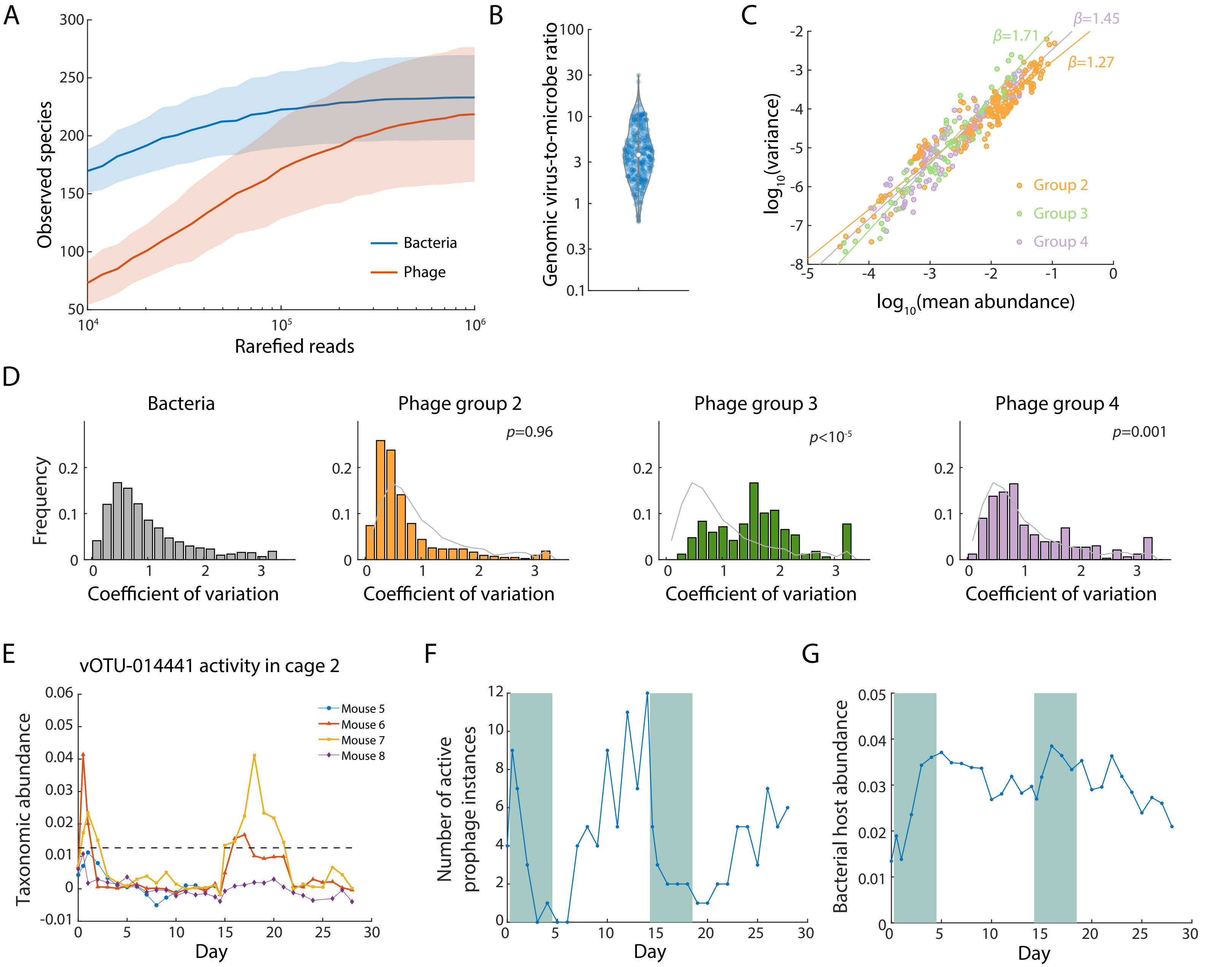
